# Neural Circuit Underlying Individual differences in Visual Escape Habituation

**DOI:** 10.1101/2024.09.05.611366

**Authors:** Xuemei Liu, Juan Lai, Chuanliang Han, Kang Huang, Qin Yang, Yuanming Liu, Xutao Zhu, Pengfei Wei, Liming Tan, Fuqiang Xu, Liping Wang

## Abstract

Emotions, like fear, are internal states enabling organisms to effectively confront environmental threats. When repeatedly exposed to predators, individuals show divergent adaptive responses. However, the neural circuit mechanisms underlying individual differences in to repeated threats remain largely unknown. Here, we identify two distinct types of visual escape—consistent escape (T1) and rapid habituation (T2) — with unambiguous arousal states to repetitive threat stimuli. We systematically investigate distinct pathways originating from the superior colliculus (SC) and insula that project to the basolateral amygdala (BLA), with relay stations in the mediodorsal thalamus (MD) and ventral tegmental area (VTA), mediating T1 and T2 behavioral types. Additionally, we identify the MD as a critical hub integrating SC and insula inputs, projecting to the BLA and contributing to reduced arousal and attenuated defensive behaviors against looming stimuli. Our findings offer significant insights into the mechanisms of internal states, arousal modulation, and behavioral adaptability.

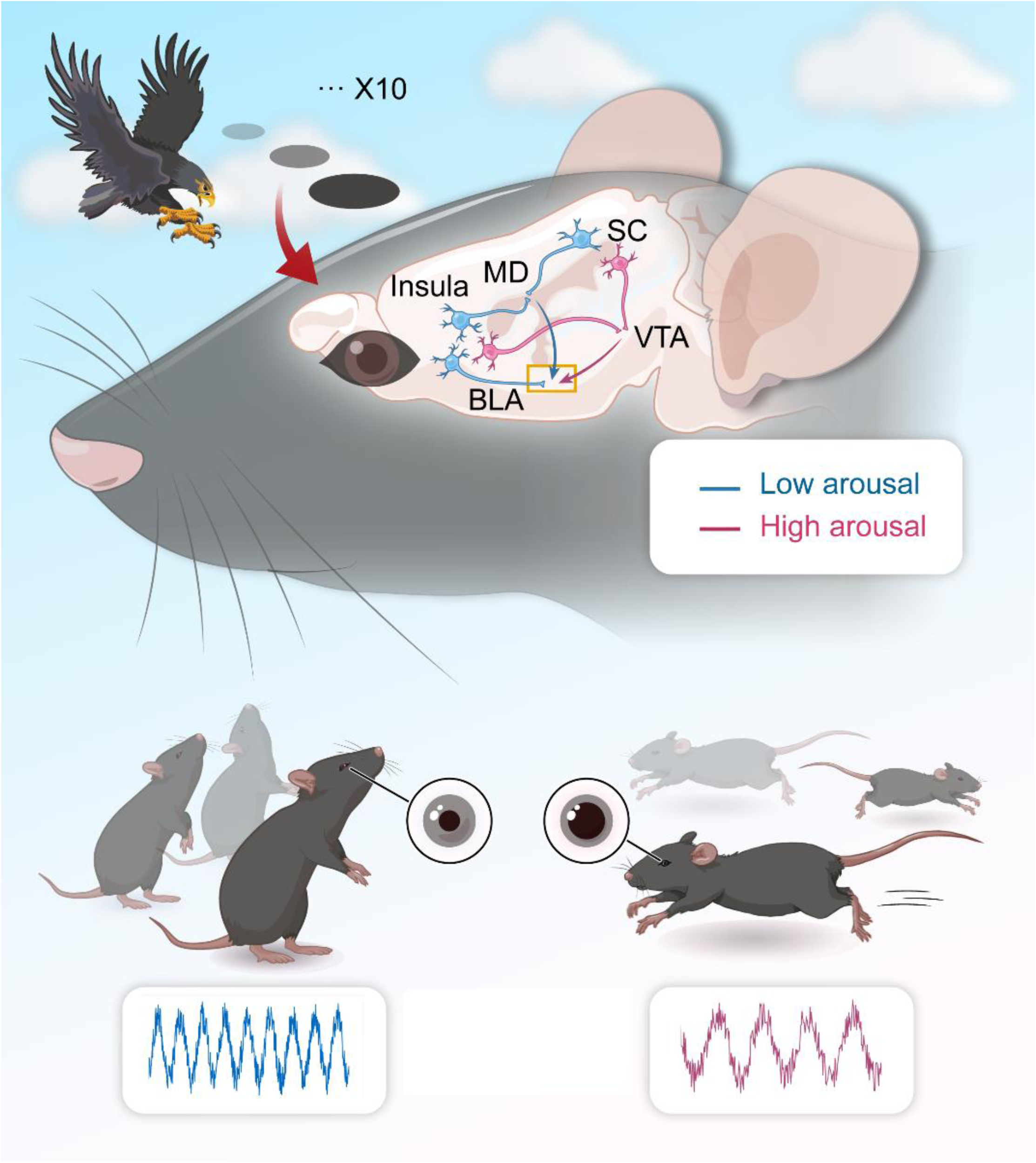
Graph Abstract.

**In brief:** We identify two distinct visual escape behaviors—consistent escape (T1) and rapid habituation (T2)—linked to unique SC and insula pathways projecting to the BLA via MD and VTA. The MD acts as a hub integrating sensory inputs to modulate arousal and defensive responses. These findings advance our understanding of the neural mechanisms driving internal states, arousal modulation, and behavioral adaptability.

**Highlights:**

● We identify two distinct visual escape behaviors with unambiguous arousal states—consistent escape (T1) and rapid habituation (T2) —to repeated looming stimuli.
● The SC-VTA-BLA pathway mediated T1 behavioral type, while the SC-MD-BLA pathway mediated T2 type.
● MD as a central hub integrating SC and insula inputs to modulate arousal and defensive behavioral adaptation.

## Introduction

Emotional responses, including fear behaviors, are hallmark features of internal states, hardwired in the brain to facilitate threat avoidance and coping ^1–5^. Arousal states typically heighten the probability of exhibiting specific behaviors or influence the behavioral choice^4,6–8^. Notably, an animal’s capacity to quickly adapt to external challenges is driven by the biological significance of repeated predator encounters compared to singular events. Individuals, across various species, exposed to recurring threats may express two distinct coping strategies habituation and sensitization dependent on sensory inputs, internal states, and previous experiences^9–12^. The neural circuits underpinning individual internal state regulation and habituation in response to repeated predator threats remain largely elusive.

Visual looming stimuli (LS), which simulate predator approach, elicit strong innate defensive responses across various species^13–18^, along with habituation behaviors upon repeated exposure^19–23^. Effective adaptive defense mechanisms necessitate a heightened state of awareness, an optimal level of arousal, and a focus on visually salient and biologically relevant stimuli. Previous research indicates that the magnitude and intensity of innate escape responses are influenced by external environmental factors and the organism’s internal state^24–31^. Despite these insights, the neural circuitry underlying the modulation of arousal levels and habituation in response to repeated LS remains inadequately understood.

The superior colliculus (SC), a retina-receipt brainstem that receive inputs from multiple sensory modalities and controls orienting, saccadic eye movement and defensive behaviors^32^. It is instrumental in capturing and allocating attention, directing both eye movements and covert attention toward visually salient stimuli^33–36^. The “subcortical route to the amygdala,” also known as the “innate alarm system,” originates from the SC and bypasses the cortex, serving as a rapid and efficient shortcut for threat avoidance^36^. The ventral tegmental area (VTA) is a SC-originating pathway, which is a crucial component in motivation control, value evaluation, and salience detection following an aversive stimulus^37–39^ and even innate defensive responses^40,41^.

Consequently, the SC-VTA-amygdala pathway is well-suited for facilitating fear learning and mediating habituation or dishabituation effects^40^. Additionally, the mediodorsal thalamus (MD), a downstream target of the SC, is involved in processing visual salience and arousal levels induced by sensory stimuli^42–44^. The SC-MD-amygdala pathway has been identified as mediating persistent fear attenuation, contributing to the underlying neurobiology that modulates visual-attentional processes^45^.

The insular cortex is pivotal in detection internal states, risk assessment, decision-making, and promoting bodily and self-awareness^46–50^. Previous research demonstrates that the insula processes fear-related emotions by modulating the amygdala, a key node in subcortical pathways^51–53^. However, the coordination between insula cortex and subcortical pathways that modulate defensive arousal and habituation to repeated predator exposures via the amygdala remains underexplored. Surprisingly, we found that approximately one-third of the mice (T2) exhibited rapid habituation characterized by low arousal (as indicated by decreased pupil size) but high attention levels (demonstrated by elevated rearing frequency) in response to repeated LS. Conversely, the remaining two-thirds of the mice (T1) consistently exhibited escape responses with corresponding arousal and rearing frequencies. We identified parallel pathways originating from the SC and insula that project to the amygdala, using relay stations in the MD and VTA, which separately mediate the T1 and T2 behavioral responses. Our study systematically explores the neural circuit mechanisms underlying distinct behavioral choices when facing repeated threats, shedding light on their internal states, arousal, and behavioral adaptability.

## RESULTS

### Individual Variability in Escape Habituation to Repeated Looming Stimuli

Previous studies have established that looming stimuli (LS) reliably elicit innate escape behaviors. To explore the escape habituation to repeated LS, we exposed 52 wildtype (WT) adult male mice to 10 LS trials per session, with each trial lasting 5.5 seconds and inter-stimulus intervals (ISIs) of no less than 2 minutes (Figure 1A). Notably, our findings revealed individual differences in escape behavior. Only one mouse (∼2%) exhibited no response, while the remaining 51 mice were categorized into two distinct response types. Mice with an average response latency shorter than 5.5 seconds were designated as “consistent escape” (Type Ⅰ, T1, n=35, 67.31%), while those with latencies exceeding 5.5 seconds were identified as exhibiting “rapid habituation” (Type Ⅱ, T2, n=16, 30.77%) (Figure 1B). Compared to the T1 group, mice in the T2 group had a significantly lower average escape proportion (35.63% vs. 98.57) (Figure 1C), and exhibited longer average response latency (17.78±1.10 s vs 1.56±1.10 s), longer return time (19.86±0.31 s vs 2.91±1.09 s), and spent less time in the nest (24.68±0.59 s vs 72.96±1.15 s) (Figure 1F,G).

**Figure 1.**
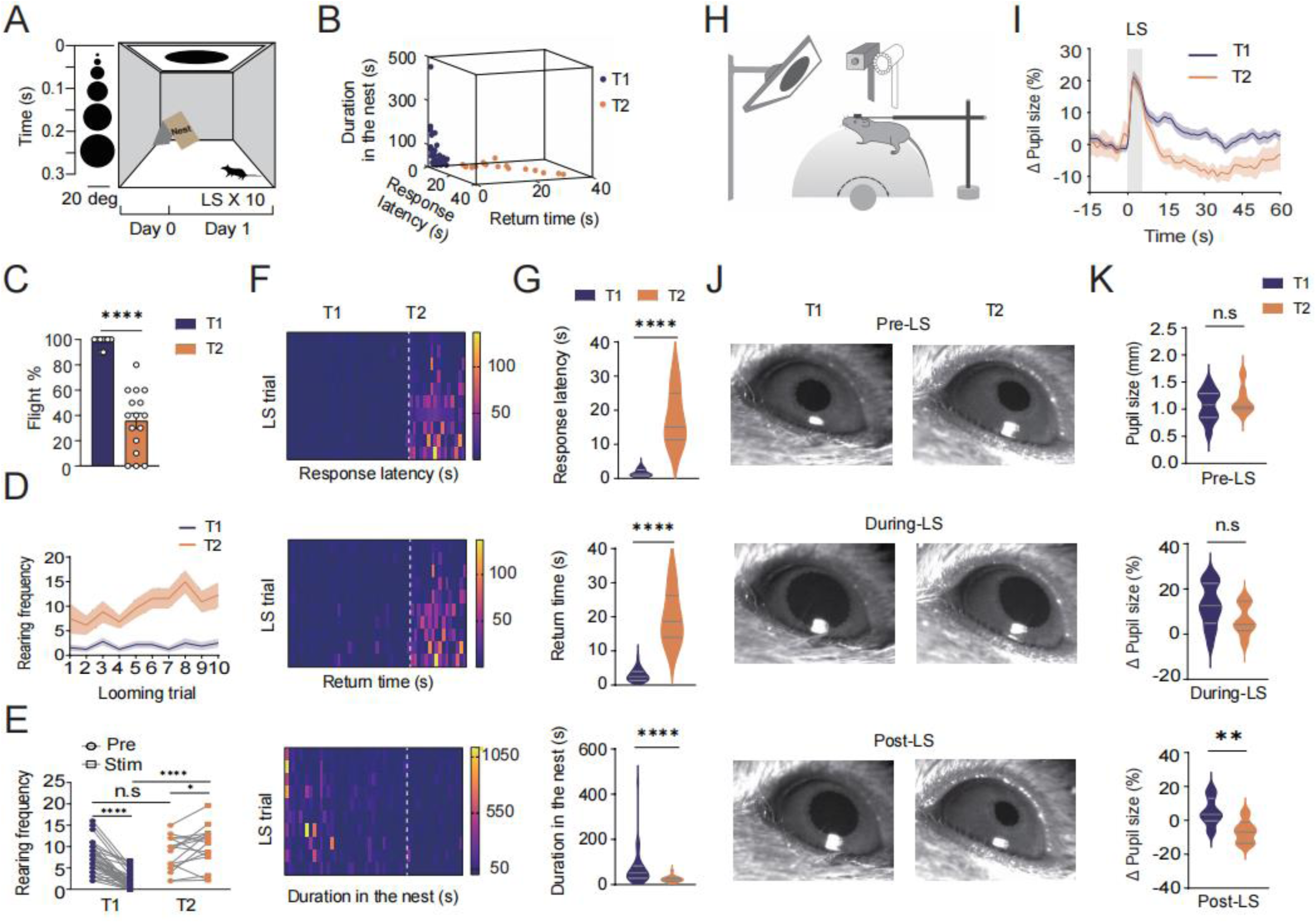
Individual Variability in Escape Habituation to Repeated Looming Stimuli. **(A)** Schematic paradigm and timeline of 10 blocks of LS. A dark disk expanding from 2 to 20°in 5.5 s. Each ISIs was random within a 2-min period. **(B)** 3D scatter plot of escape behavior parameters (response latency, time to return to nest and duration in nest) of 51 male mice categorized as one of two subgroups. T1, Type1, “low habituation” group, T2, Type 2, “high habituation” group. **(C)** The average proportion of trials in which escape occurred in T1 and T2 mice. **(D)** Comparison of rearing frequency in each trial in T1 and T2 mice. **(E)** Comparison of rearing frequency pre-LS (1 min before first LS) and during-LS in T1 and T2 mice. (Two-tailed unpaired t test, p=0.3750, ****P<0.0001). **(F)** Heatmap showing escape behavior parameters in each trial of the 51 mice. **(G)** Bar graph showing the average 1) response latency (Two-tailed unpaired t test, ****p<0.0001), 2) return time (Two-tailed unpaired t test, ****p<0.0001) and 3) duration in nest (Two-tailed unpaired t test, ***p=0.0002) of T1 and T2 mice responding to 10 blocks of LS. **(H)** Schematic of pupillometry recording paradigm with LS in head-fixed awake behaving mice on a ball treadmill. **(I)** Graph showing the percentage of pupil-size change 15 s before LS and 60 s after LS in T1 and T2 mice. **(J)** Representative image showing pupil diameter 5 s before LS, 5.5 s during LS and 10s after LS. **(K)** Pupil size 5 s before LS, 5.5 s during LS and 10 s after LS.

Analysis of average scores over trials revealed that the T2 group showed increased latency and time to return to the nest across initial, middle, and final trial phases (Figure S1).

Non-selective attention (NSA), characterized by scanning, orienting, and detecting stimuli, correlates with rearing frequency in novel settings ^54–56^. The T2 group consistently exhibited higher rearing frequency (RF) across all trials compared to the T1 group (Figure 1D). Remarkably, in the T2 group, RF significantly increased following LS onset, in contrast to a decrease observed in the T1 group. No RF differences were noted between T1 and T2 groups before LS onset, indicating stimulus-specific RF patterns (Figure 1E). To assess arousal state, we performed pupillometry on head-fixed, awake mice placed on a ball treadmill (Figure 1H). Pupil size, measured 10 seconds post-LS onset, was significantly larger in the T1 group compared to the T2 group (Figure 1 IJK). No significant pupil size differences were observed between groups prior to or during LS exposure. Collectively, our results demonstrate that the T2 group, characterized by higher habituation, exhibits enhanced stimuli-evoked NSA and reduced arousal in response to repeated LS presentations.

### Divergent Superior Colliculus Pathways Modulate Arousal and Escape Behavior

To elucidate the role of SC neurons activated during habituation to LS, we employed the FosTRAP2 technique ^57,58^. On day 1, AAV5-DIO-EGFP was injected unilaterally into the SC of FosTRAP2 mice. On day 3, FosFRAP2 mice were injected with tamoxifen and exposed to repeated LS to activate TRAPed cells. On day 24, FosFRAP2 mice were exposed to LS again before perfusion (Figure 2A). Our results showed a higher density of EGFP+ SC neurons in the T1 group compared to the T2 group, suggesting the SC’s involvement in LS habituation and attenuation of innate escape behavior (Figure 2B-C).

**Figure 2.**
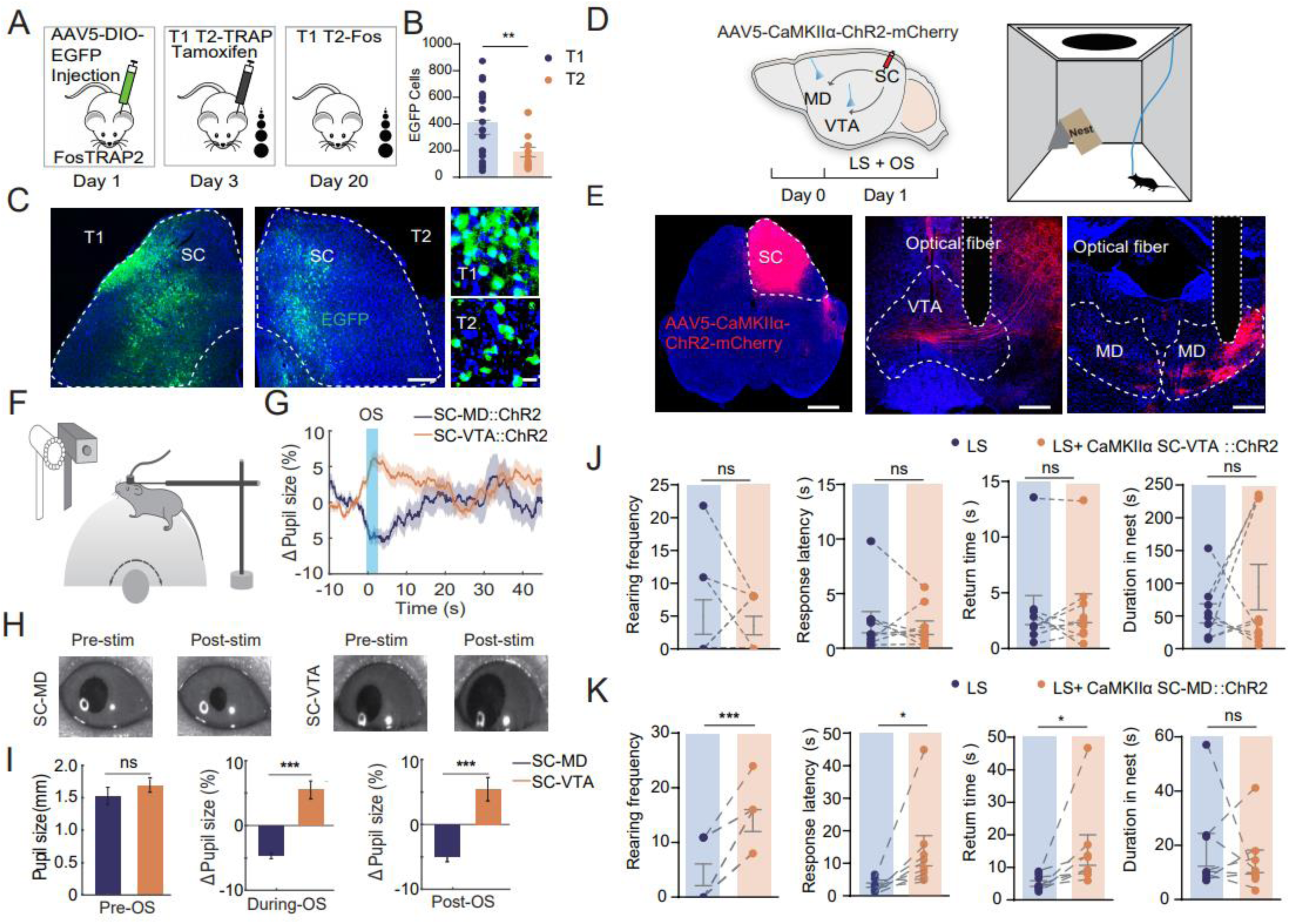
Divergent Superior Colliculus Pathways Modulate Arousal and Escape Behavior. (**A**) Schematic showing the FosTrap2 procedure. (**B**) The number of EGFP+ neurons in the SC in T1 and T2 mice (Two-tailed unpaired t test, **p=0.0032, n=22 slices from 4mice in T1, n=15 slices from 4 mice in T2). (**C**) Looming-associated EGFP+ neuronal activity expressed in the SC in T1 and T2 mice. (**D**) Schematics showing the OS (optogenetic stimuli) paradigm (*left*) and the experimental procedure with LS (looming stimuli, *right*). (**E**) Injection of AAV5-CaMKIIα-ChR2-mcherry in the IDSC and fiber implantation in the VTA and MD. Optical fiber positions are outlined using white dotted lines. Blue, DAPI, scale bar=20μm. (**F**) Schematic paradigm of pupillometry recordings with OS in head-fixed awake behaving mice on a ball treadmill. (**G**) Graph showing the percentage of change in pupil size 10 s before OS and 45 s after OS of the VTA and the MD. (**H**) Representative example of pupil diameter 5 s before OS, and 10 s after OS. (**I**) Bar graph showing the change in pupil size 10 s before OS (two sample t test, p=0.35), 2.5 s during OS (two sample t test, p<0.001) and 2.5 s after OS (two sample t test, p<0.001). (**J**) Measurements were taken during OS (10 trials/mouse) and LS of CaMKIIα_SC-VTA_:: ChR2 mice. Bar graphs showing 1) rearing frequency (two-tailed paired t test, p>0.05), 2) response latency (two-tailed paired t test, p>0.05), 3) return time (two-tailed paired t test, p>0.05), and 4) duration in nest (two-tailed paired t test, p>0.05). (**K**) Measurements were taken during OS (10 trials/mouse) and LS of CaMKIIα_SC-MD_:: ChR2 mice. Bar graphs showing 1) rearing frequency (two-tailed paired t test, ***p=0.0004), 2) response latency (two-tailed paired t test, *p=0.042), 3) return time (two-tailed paired t test, *p=0.0497), and 4) duration in nest (two-tailed paired t test, p>0.05).

We investigated whether the SC-VTA and SC-MD pathways differentially affect arousal during LS habituation. Using AAV5-CaMKIIα-ChR2-mcherry injections into the SC and optic fiber implantation into the MD and VTA (Figure 2D-E). we performed pupillometry on head-fixed mice. Optogenetic stimulation (473 nm, 2.5 s, 20 Hz, 50 pulses) altered pupil size during LS presentation. SC-MD activation led to pupil constriction, while SC-VTA activation resulted in pupil dilation (Figure 2F-G). Before optogenetic stimulation (OS), there was no significant difference in pupil size between the groups. However, the mean pupil size during and after OS was significantly larger in the SC-VTA group than in the SC-MD group (Figure 2H-I).

Considering the larger pupil size in the T1 group and following SC-VTA activation, we hypothesized that repeated SC-VTA stimulation would elicit T1-like behavior. Indeed, repeated SC-VTA OS resulted in stable escape behavior in an open field with shelter, mimicking T1 group responses (Figure S2).

Further investigations assessed the SC pathways’ roles via ChR2-mcherry expression in SC neurons. SC-MD OS increased rearing frequency, response latency, and return time but SC-VTA OS did not, indicating SC-MD OS disrupts looming-induced escape behaviors. Thus, SC-MD activation decreased arousal and increased NSA, impairing innate escape, whereas SC-VTA activation increased arousal without affecting NSA or escape behavior.

To trace SC projections, we injected rabies virus (RV) conjugated with dsRed into the VTA and RV-EGFP into the MD, quantifying labeled neurons across the brain. Among retrogradely labeled neurons, 51% in the SC were from the MD, and 63% from the VTA, with 14.4% co-labeled from both sites (Figure S3), indicating SC collaterals project to both MD and VTA.

To rule out collateral effects, we blocked SC action potential backpropagation using 0.3 μl of 4% bupivacaine ^59^. Bupivacaine had no impact on behavior change during either SC-VTA or SC-MD pathway activation in an open field with shelter paradigm (Figure S3), suggesting parallel processing between SC pathways modulates variability in innate escape responses.

### VTA-projecting and MD-projecting SC neurons functionally target the BLA

The amygdala is pivotal for both conditioned and innate fear responses, with the basolateral amygdala (BLA) playing a central role in encoding threat-related information ^3,60^. We investigated whether SC neurons projecting to the VTA and the MD converge on the BLA. Using AAV1-mediated anterograde transsynaptic tagging ^61^, we injected AAV1-Cre into the SC, followed by injections of AAV5-DIO-EYFP into the MD and AAV5-DIO-mCherry into the VTA (Figure 3A). Cre expression revealed overlapping axonal projections in the BLA, indicating that both VTA-projecting and MD-projecting SC neurons innervate a common region within the BLA (Figure 3B). Fluorescence density analysis showed that VTA-projecting SC neurons primarily innervated the medial BLA, while MD-projecting neurons targeted the anterior and posterior parts (Figure 3C).

**Figure 3.**
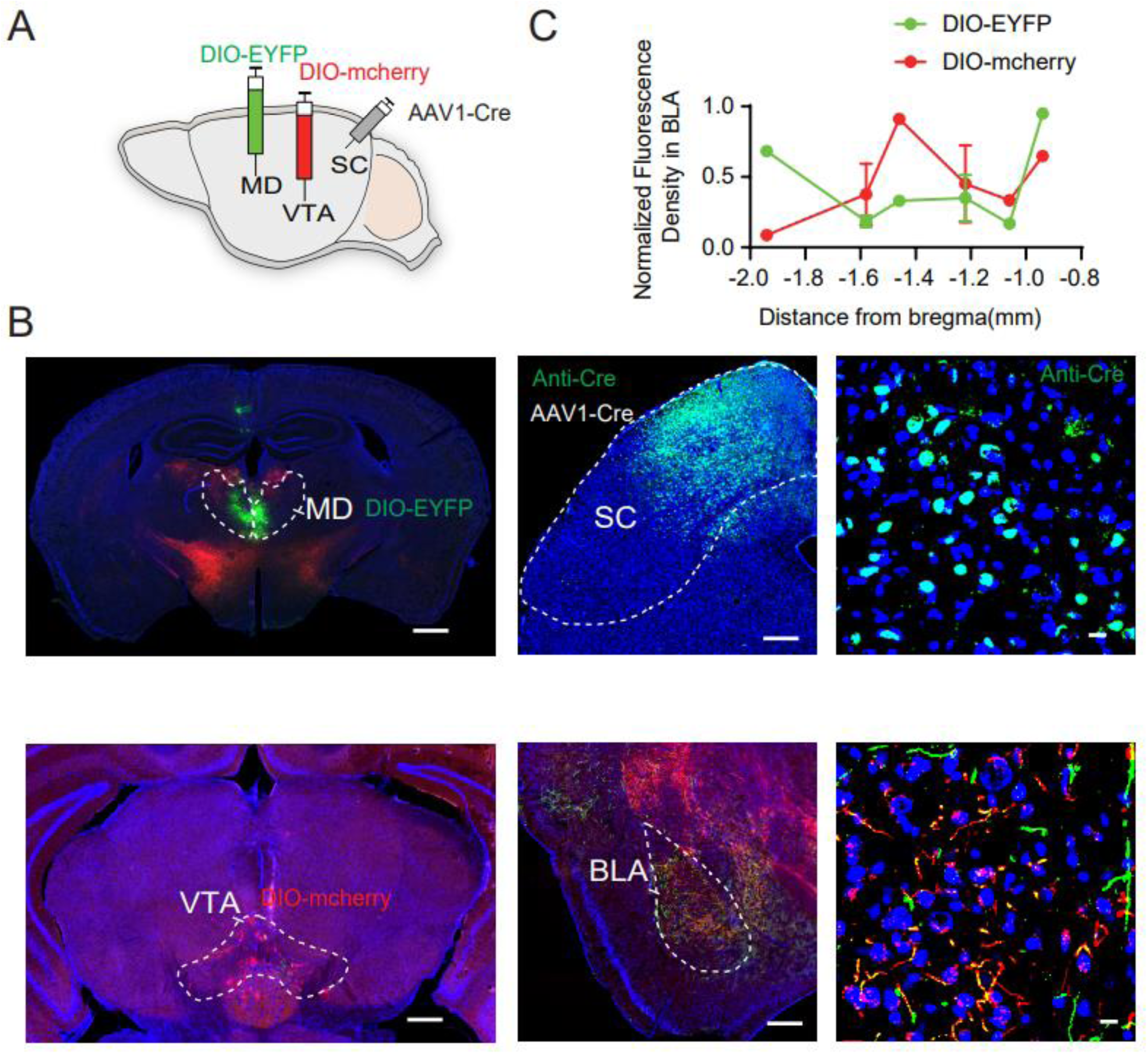
VTA-projecting and MD-projecting SC neurons target the BLA. **(A)** Schematic showing injections of AAV1-Cre into the SC and AAV5-DIO-mchery and AAV5-DIO-EYFP into the VTA and the MD. **(B)** Representative images showing AAV1-Cre virus injection and Cre immunopositive expression in the SC, AAV5-DIO-EYFP in the MD, AAV5-DIO-mcherry in the VTA and fiber terminals in the BLA. Scale bar= 250μm (left), 100μm (middle), 10μm (right). **(C)** Normalized fluorescence density of fibers labeled with mcherry and eYFP in the BLA.

### SC Pathways Modulate Connectivity and Oscillations in the BLA

To explore this functional connectivity further, we conducted in vivo multi-channel recordings in the BLA and pupillometry during selective optogenetic activation of SC-VTA and SC-MD pathways (Figure 4A). OS of the SC-VTA pathway caused an increase in pupil size, while SC-MD activation led to a decrease (Figure 4B). Neuronal firing rates within the BLA significantly changed following the activation of each pathway; SC-MD activation affected 11% of neurons (9% excitation, 2% inhibition), and SC-VTA activation affected 9% (7% excitation, 2% inhibition) (Figure S4,S5).

**Figure 4.**
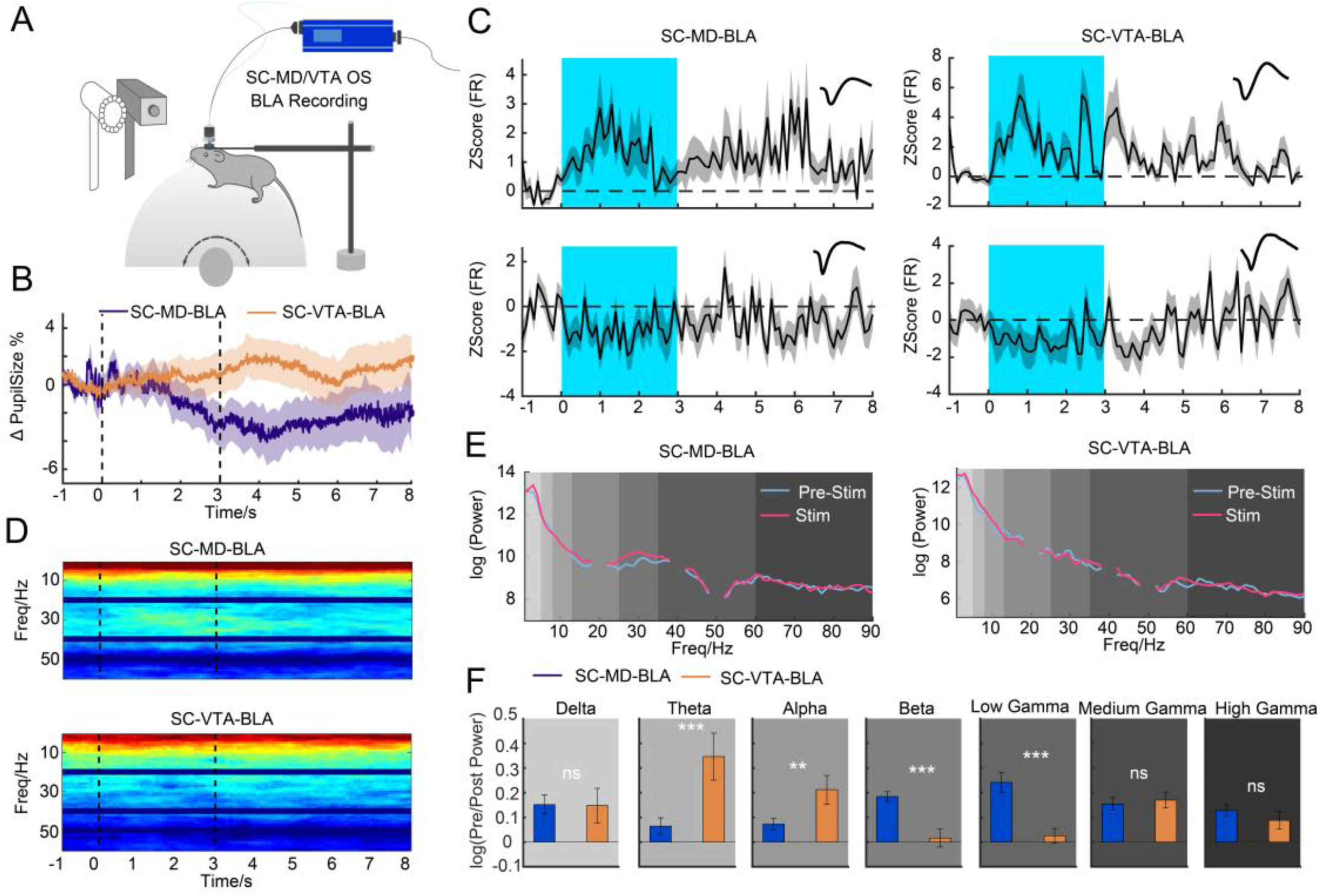
SC Pathways Modulate Connectivity and Oscillations in the BLA. **(A)** Schematic showing electrophysiological recording of VTA-projecting and MD-projecting SC neurons target BLA, and an eye tracker to monitor the pupil size. **(B)** Graph showing normalized pupil size during optogenetic stimulation (inside the two black dashed lines) comparing VTA-projecting SC neurons (yellow) and MD-projecting SC neurons (blue). **(C)** Graph showing the normalized firing rate (z-scores) of the neurons recorded in the BLA during optogenetic stimulation. Examples show two types of neuron (excitation and inhibition; spike waveforms are shown above) that were modulated by optogenetic stimulation (blue region, optogenetic stimulation). **(D)** LFP spectrograms recorded in the BLA during optogenetic stimulation (inside the two black lines). *Top* example from VTA-projecting SC neurons; *Bottom* example from MD-projecting SC neurons. **(E)** Power spectrums in BLA comparing before (light blue line) and during (pink line) optogenetic stimulation for VTA-projecting and MD-projecting SC neurons. **(F)** Bar graph showing the comparison between LFP power in different frequency bands before, during and after optogenetic stimulation (two sample t test, delta, p=0.95; theta, p<0.001; alpha, p=0.0029; beta, p<0.001; low gamma, p<0.001; medium gamma, p=0.66; high gamma, p=0.30).

The analysis of local field potential (LFP) power spectra in the BLA revealed distinct oscillatory patterns associated with each pathway. SC-VTA activation led to increased power in low-frequency bands (theta and alpha), whereas SC-MD activation enhanced power in higher frequency bands (beta and low gamma) (Figure 4D-F). These findings underscore the distinct oscillatory characteristics modulated by SC-MD and SC-VTA pathways, highlighting their unique roles in amygdala circuitry and fear-related behaviors.

### Distinct Arousal and Defense Modulation by Divergent Insula Cortex Pathways

The insula cortex, a key integration center for sensory feedback and autonomic arousal ^46^, projects to both the MD and the VTA, as confirmed by our previous tracing studies (Figure S3). To assess whether the insula-MD and insula-VTA pathways modulate arousal differently, we used optogenetic manipulation and pupillometry (Figure 5A). AAV5-CaMKIIα-ChR2-mCherry was injected into the insula cortex, with optical fibers placed in the MD and VTA (Figure 5D). OS of MD terminals significantly decreased pupil size, while OS of VTA terminals increased pupil size (Figure 5B, 5E-F). No significant differences in baseline pupil size were observed between groups prior to OS (Figure 5E-F). During and post-stimulation, pupil size was smaller in the insula-MD group than in the insula-VTA group, indicating distinct effects on arousal (Figure 5E-F).

**Figure 5.**
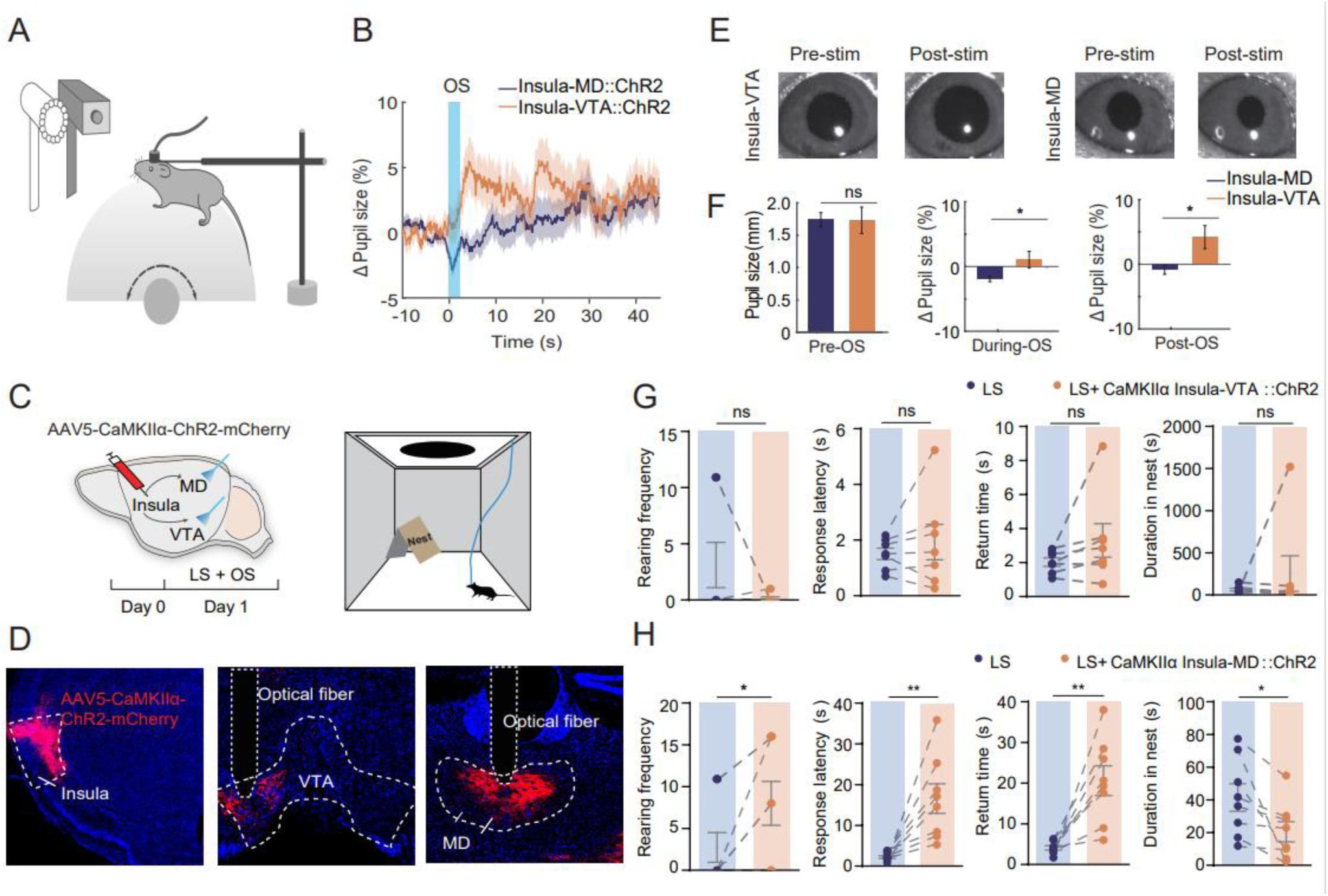
Divergent Insula Cortex Pathways modulate arousal and innate escape behavior. (**A**) Schematic showing the pupillometry recording paradigm. (**B**) Graph shows the percentage of pupil size change at 10 s before OS (blue shaded bar), 2.5 s during OS and 45 s after OS. (**C**) Schematic showing the OS paradigm and the LS set-up. (**D**) Schematic showing injection of AAV-CaMKIIα-ChR2-mcherry into the insula cortex and fiber implantation in the VTA and the MD. Blue, DAPI, scale bar=20 μm. (**E**) Representative example of pupil diameter size 5 s before and 10 s after OS of either the insula-MD or the insula-VTA pathway. (**F**) Bar graph showing pupil size change 10 s before OS (Two sample t test, p=0.97), 2.5 s during OS (Two sample t test, p=0.0106) and 2.5 s after OS (Two sample t test, p=0.0025). (**G**) Measurements were taken during OS (10 trials/mouse) and LS of CaMKIIα_insula-VTA_:: ChR2 mice. Bar graphs showing 1) rearing frequency (two-tailed paired t test, p>0.05), 2) response latency (two-tailed paired t test, p>0.05), 3) return time (two-tailed paired t test, p>0.05), and 4) duration in nest (two-tailed paired t test, p>0.05). Measurements were taken during OS (10 trials/mouse) and LS of CaMKIIα_insula-MD_:: ChR2 mice. Bar graph showing 1) rearing frequency (two-tailed paired t test, *p=0.0306), 2) response latency (two-tailed paired t test, **p=0.0067), 3) return time (two-tailed paired t test, **p=0.0033), and 4) duration in nest (Two-tailed paired t test, *p=0.0315).

We explored the involvement of these pathways during LS presentations by testing mice with ChR2-mCherry in insula cortex neurons, Activation of the insula-MD pathway increased rearing frequency, response latency, and return time, while reducing time spent in the nest. However, insula-VTA activation did not affect LS-evoked behaviors. Thus, insula-MD activation decreased arousal levels and increased non-selective attention (NSA), diminishing LS-induced escape responses, whereas insula-VTA activation heightened arousal without influencing escape (Figure 5C-D, 5G-H). To further investigate functional connectivity, we conducted in vivo multi-channel recordings in the BLA and pupillometry during insula-MD pathway activation (Figure 6A). Insula-MD stimulation decreased pupil size (Figure 6B) and altered neuronal firing rates in the BLA, with 30% of neurons responding (8% excitation, 22% inhibition) (Figure S6); Although LFP power spectra in the BLA were analyzed, insula-MD activation did not significantly alter powers across frequency bands (Figure 6D-F).

**Figure 6.**
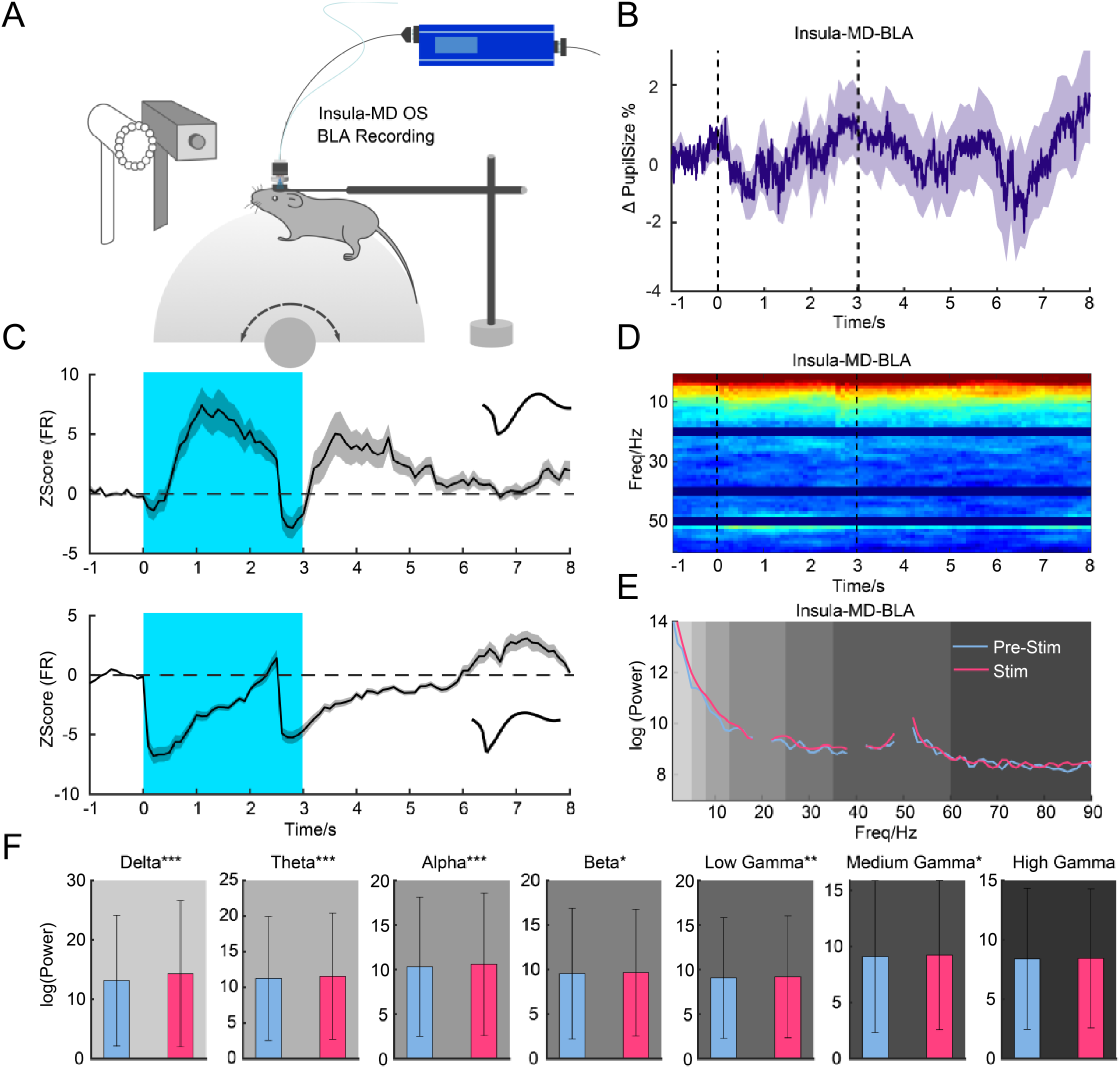
Insula-MD pathway modulate oscillations in the BLA. **(A)** Schematic showing electrophysiological recording in the BLA following OS of VTA-projecting and MD-projecting insula neurons. **(B)** Graph showing normalized pupil size during OS (inside the two black dashed lines). **(C)** Graph showing the firing rate of neurons recorded in the BLA during OS. Examples show two types of neuron (excitation and inhibition; spike waveforms are shown above) that were modulated by optogenetic stimulation (blue shaded region). **(D)** Time course of LFP power in alpha and gamma band (optical stimulation starts and ends between black dotted lines). **(E)** Averaged LFP spectrograms recorded in the BLA during OS (inside the two black lines) **(F)** Bar graph showing the comparison between the LFP power in different frequency bands before, during and after optogenetic stimulation (no significant difference)

We examined whether the insula-BLA pathway impacts arousal using optogenetic manipulation and pupillometry (Figure 7). We performed optogenetic manipulation combined with pupillometry recording as described above (Figure 7A). Following ChR2-mCherry injections into the insula cortex and fiber placement in the BLA, OS reduced pupil size (Figure 7 B-D). Insula-BLA activation increased rearing frequency, response latency, and return time, suggesting reduced LS-induced escape behaviors. In further recordings (Figure 8A), insula-BLA activation affected 42% of BLA neurons (40% excitation, 2% inhibition, Figure S7) and decreased pupil size (Figure 8B-E); LFP power analyses showed no significant changes, contrasting with SC-MD activation and suggesting a more pronounced role for subcortical versus cortical pathways in modulating escape responses (Figure 8D-F).

**Figure 7.**
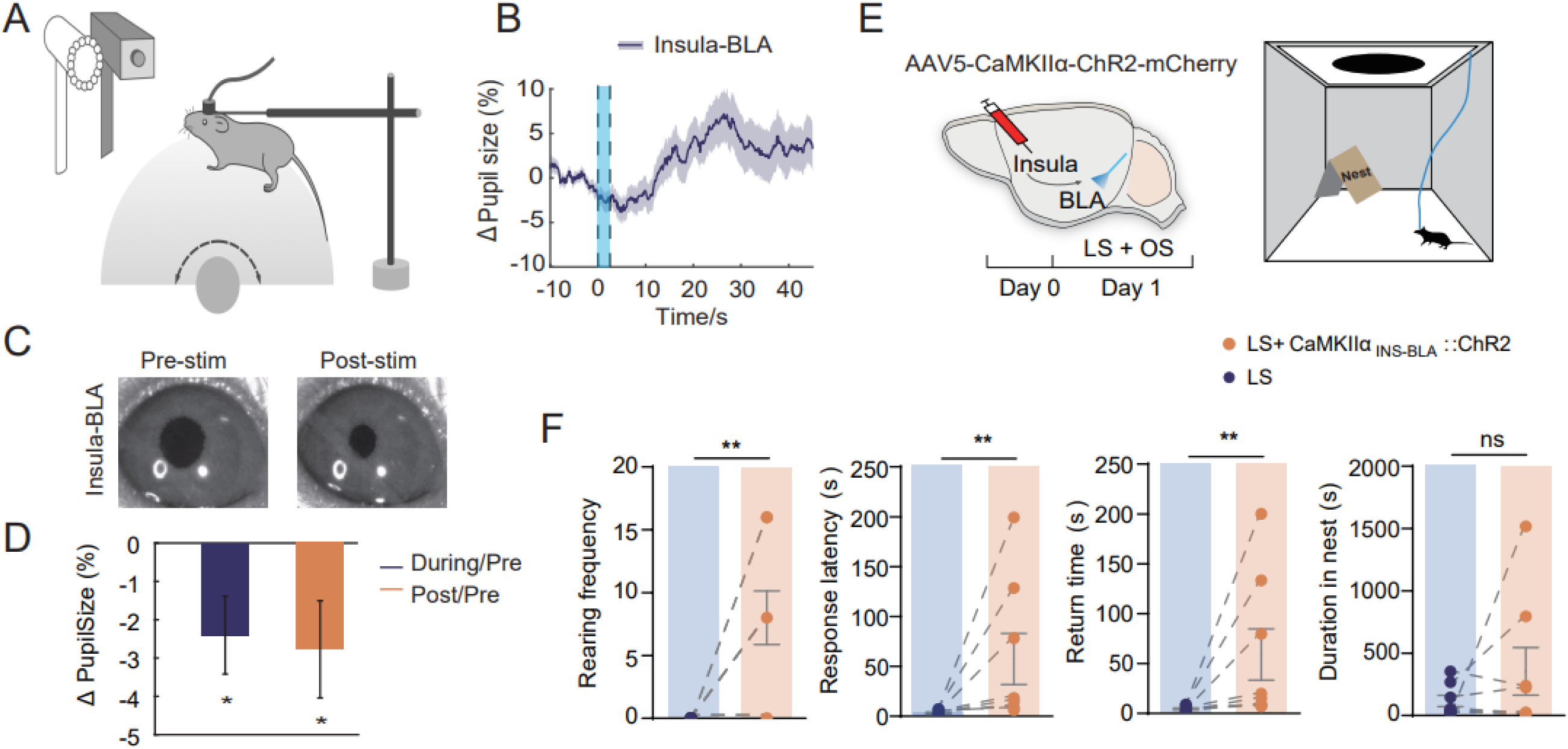
Influence of Insula-BLA Pathway on Arousal and Escape Behavior. **(A)** Schematic paradigm showing the pupillometry recording set-up. **(B)** Graph showing the percentage of change in pupil size 10 s before OS, 2.5 s during OS, and 45 s after OS. **(C)** Representative image showing pupil diameter 10 s before OS, 2.5 s during OS and 2.5 s after OS. **(D)** Bar graph showing change in pupil size During/Pre OS (Two-tailed paired t test, *p=0.0195) and Post/Pre OS (Two-tailed paired t test, *p=0.0294). **(E)** Schematic showing the OS paradigm and the LS set-up (**H**) Measurements were taken during OS (10 trials/mouse) and LS of CaMKIIα_insula-MD_:: ChR2 mice. Bar graphs showing 1) rearing frequency (two-tailed paired t test, **p=0.0072), 2) response latency (Wilcoxon matched-pairs signed rank test, **p=0.0078), 3) return time (Wilcoxon matched-pairs signed rank test, **p=0.0078), and 4) duration in nest (two-tailed paired t test, p>0.05).

**Figure 8.**
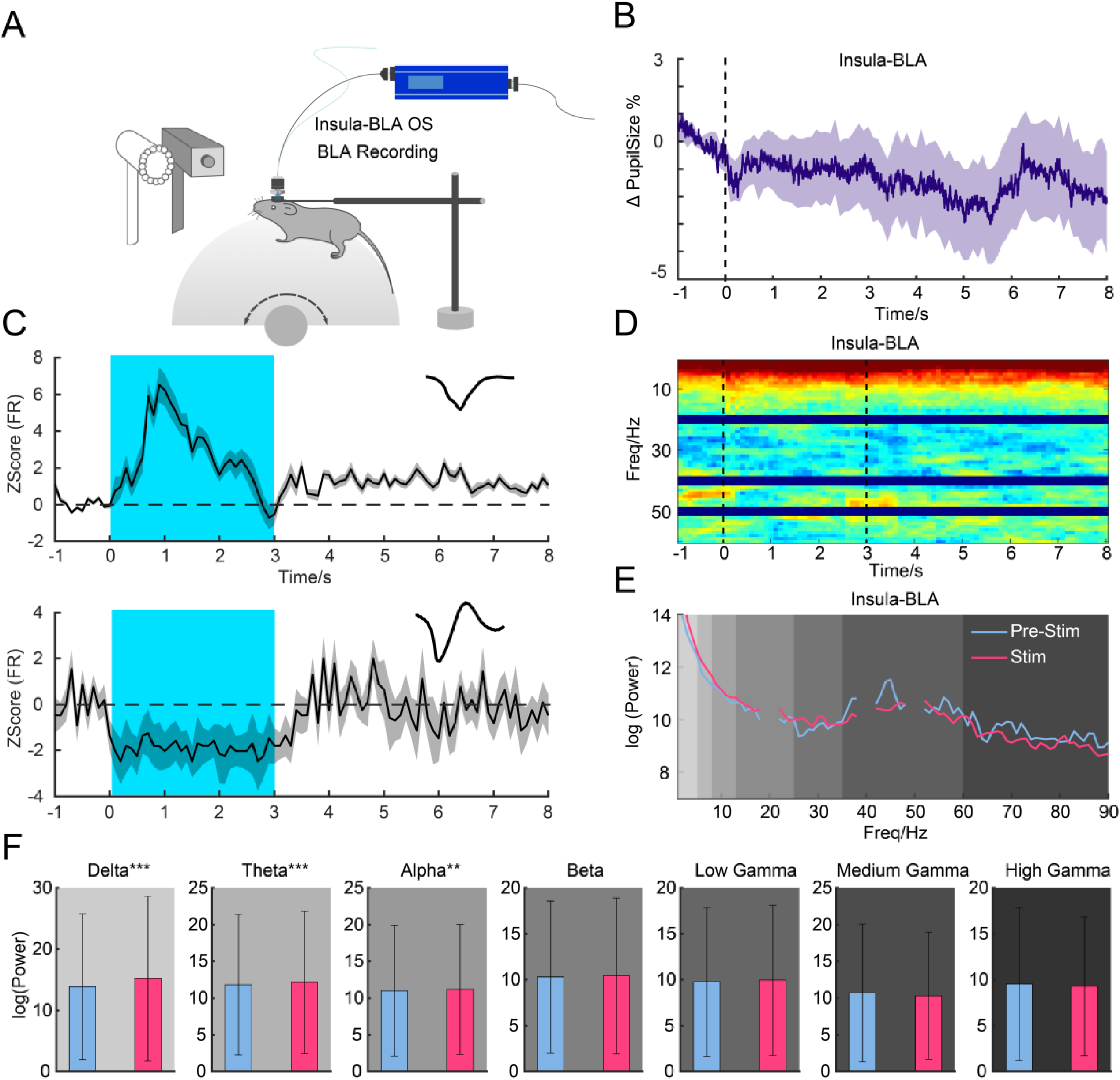
Insula-BLA pathway modulate oscillation in the BLA. **(A)** Schematic showing electrophysiological recording of neurons in the BLA during OS of the insula BLA pathway. **(B)** Graph showing normalized pupil size during OP (inside the two black dashed lines) of the insula-BLA pathway. **(C)** Firing rate of the neurons recorded in BLA during OS. Examples show two types of neuron (excitation and inhibition; spike waveforms are shown above) that were modulated by optogenetic stimulation (blue region, optogenetic stimulation). **(D)** Time course of LFP power in alpha and gamma band (OS inside the two black lines). **(E)** Averaged spectrogram of LFP recorded in the BLA during OS (inside the two black lines). **(F)** Bar graph showing the comparison between the LFP power in different frequency bands before, during and after optogenetic stimulation (no significant difference)

## Discussion

In the study, we found that rodents exhibit two types of behavioral responses to repeated light stimulation (LS): “consistent escape” (T1) and “rapid habituation” (T2), each characterized by distinct pupil sizes and rearing frequencies. Additionally, we identified divergent pathways originating from the SC and the insula, which project to BLA and contribute to different levels of defensive arousal and habituation responses to repeated LS. Furthermore, the state of defensive arousal and the manipulation of separate circuits play distinct roles in regulating the power of LFP in the BLA. This research enhances our understanding of how internal states, such as arousal and fear, are modulated at the neural level, and highlights the complexity of these processes.

Individual differences are crucial as they supply the raw materials for natural selection, as evidenced by the variability in individual adaptations to repeated predator encounters^62,63^. The behavioral adaptation need optimizing perception and attention by conserving cognitive resources, enhancing biological salience detection thereby enable threat detection and resilience ^52,64^. The SC’s well-documented functions in detecting and directing attention to salient visual stimuli align with its involvement in both T1 and T2 behaviors. In T1, consistent escape behavior is driven by sustained activation of the SC-VTA-amygdala pathway, maintaining the high arousal needed for continuous defensive responses, which, despite its energy cost, this sensitization-like response enhances survival probability in environments with unpredictable threats^65^. This pathway’s role in reinforcement fear learning and mediating potential sensitization or dishabituation effects is consistent with its established contributions to innate defensive responses and motivational control^32,37^.

Conversely, T2 behavior involves rapid habituation via transient SC-MD-amygdala pathway or insula-MD-amygdala pathway activation, swiftly reducing arousal and defensive responses. This rapid habituation aligns with previous studies in larval zebrafish, crabs, and other species ^66–68^, enabling animals to disregard irrelevant stimuli and focus on biologically significant threats, thereby enhancing adaptive survival by reducing responses to non-threatening cues. The SC also influences higher-order regions by directing attention in a bottom-up manner ^69–71^. These findings underscore the role of a “visual salience network”, comprising subcortical pathways and top-down cortical visuomotor control, in coordinating visual attention towards novel, salient stimuli and unexpected threats ^49,72^.

Our study investigates the relationship between arousal and non-selective attention in rodents, as indexed by pupil size and rearing frequency, respectively. Traditionally, rearing is associated with non-selective attention, while arousal definitions vary, sometimes linked to awareness or energy states. Despite extensive research on neuromodulators affecting pupil size, the neural circuitry underpinning rearing frequency remains less understood, and their correlation unexamined. We found T1 subjects exhibit high pupil dilation and low rearing frequency, whereas T2 subjects show the opposite. This aligns with our optogenetic and electrophysiological data, suggesting distinct neural pathways from the SC and insula to the amygdala (via the VTA or MD) govern these responses. Interestingly, rearing frequency differences only emerged post-stimulus (Fig 1 E), indicating a stimulus-specific adaptation rather than innate resting state differences. This reveals a dissociation, and even a counterbalance, between arousal and non-selective attention during repeated threat exposure.

Our electrophysiological findings show differences between VTA– and MD-projecting SC neurons targeting the BLA: activation of the SC-MD circuit increases high-frequency beta (12-30 Hz) and low gamma (30-60 Hz) oscillations, while SC-VTA activation boosts lower-frequency theta (3-7 Hz) and alpha (7-12 Hz) oscillations. These patterns align with the notion that attention correlates positively with high-frequency and negatively with low-frequency oscillations. Elevated rearing frequency upon SC-MD activation suggests heightened attention due to increased beta and low gamma oscillatory power. Conversely, SC-VTA activation suggests diminished attention, correlating with enhanced engagement in escape behavior and increased lower-frequency power. Activation of the insula-MD pathway results in different oscillatory changes than SC-MD, less significantly altering attention as indicated by pupil size changes. This complexity in the insula’s regulation of the BLA suggests intricate interactions worth further investigation to understand MD and VTA functions.

This study underscores how distinct neural pathways from shared sensory inputs lead to varied behavioral outcomes based on individual internal states. By identifying the MD as a common hub projecting to the amygdala from both the SC and insula, we unveil the neural basis for modulating attention and arousal in response to threats. Our findings enhance understanding of the interplay between sensory processing and emotional states in behavior, offering a framework for future exploration of these adaptable neural circuits, central to survival in dynamic environments.

Maladaptive behaviors in response to extreme fear can lead to either heightened sensitivity to harmless stimuli (overreaction) or reduced detection and response to actual threats (underreaction) ^73–75^. In human, these responses are common in fear-related disorders such as phobias, anxiety, PTSD, and delusional disorders.

Our findings have significant implications for understanding individual variability in emotional processing and resilience. The ability to rapidly habituate to repeated threats could confer advantages in certain environments, reducing stress-related pathologies. Conversely, a consistent escape response might be advantageous in environments where threats are unpredictable and potentially lethal. Understanding the neural circuits and molecular mechanisms behind individual differences in habituation to repeated predator exposures could inform personalized treatments for neuropsychiatric conditions ^36^. By concentrating on evolutionarily conserved circuits like the SC–MD–amygdala pathway, our research establishes a physiological basis for developing potential therapies. The further detailed analysis the molecular mechanisms of the subcortical pathway could enhance the translational value of research by identifying a key target for PTSD treatment ^45,76,77^.

## Author Contributions

X.L. and L.W. designed the project, X.L., J.L., Q.Y., Y.L., X.Z. executed the experiments. X.L., C.H., J.L., K.H., analyzed the data. X.L., C.H., L.W. wrote the manuscript. X.L., L.W., P.W., L.T., F.X. revised the manuscript. L.W. and X.L. supervised the project.

## Acknowledgments

We thank Peng Cao for providing the FosTrap2 mice line (Jackson Laboratory). This work was funded by the National Natural Science Foundation of China (32230042 to L.W., 32371069 to X.L.,); Strategic Priority Research Program of CAS (XDB32030100), Financial Support for Outstanding Talents Training Fund in Shenzhen (L.W.); STI2030-Major Projects-2022ZD0211700, the Ten Thousand Talent Program, the Chang Jiang Scholars Program, Financial Support for Outstanding Talents Training Fund in Shenzhen (L.W.);Guangdong Province Basic Research Grant (2023B1515040009) and the China Postdoctoral Science Foundation (2022M723299, 2023T160666).

## STAR*METHODS

### METHOD DETAILS

#### Animal Preparation

All experimental procedures were approved by the Animal Care and Use Committees at the Shenzhen Institute of Advanced Technology (SIAT), Chinese Academy of Sciences (CAS). Adult (6– 8 week-old) male C57BL/6J (Beijing Vital River Laboratory Animal Technology Co., Ltd., Beijing, China), VGLUT2-ires-Cre (#016963) mice were used in this study. Mice were housed at 22–25 °C on a circadian cycle of 12-hour light and 12-hour dark with ad-libitum access to food and water.

#### Viral vector preparation

For optogenetic experiments, we used plasmids for AAV2/9 viruses encoding *CaMKIIa*:: hChR2 (H134R)–mCherry, *CaMKIIa*:: mCherry, *EF1α*:: DIO–hChR2 (H134R)–mCherry, *EF1α*:: DIO–mCherry, *EF1α*:: DIO–EYFP (gifts from Dr. Karl Deisseroth, Stanford University) and Retro-AAV-*EF1α-*DIO-hChR2 (H134R)– mCherry (packaged by BrainCase Co., Ltd., Shenzhen). Viral vector titers were in the range of 3-6×10^12^ genome copies per ml (gc)/mL.

For viral tracing, viral vectors RV-dG-dsRed, RV-dG-GFP, Retro-AAV-hSyn-mcherry, Retro-AAV-hSyn-eYFP, AAV1-hSyn-SV40 NLS-Cre were used (packaged by BrainCase Co.). Adeno-associated and rabies viruses were purified and concentrated to titers at approximately 3×10^12^ v.g/ml and 1×10^9^ pfu/ml, respectively.

#### Stereotaxic Surgery

Animals were anesthetized with pentobarbital (i.p., 80 mg/kg) before stereotaxic injection. The viruses were injected into the SC (AP −3.80 mm, ML ±0.8 mm, and DV −1.8 mm), the insula (AP +0.14 mm, ML ±3.75 mm, DV range between −3.75 and −3.9 mm), and the Cg (AP +0.3 mm, ML ±0.30 mm, DV −1.5 mm). Optical stimulation of terminals was conducted using a 200-µm optic fiber (NA: 0.37; NEWDOON, Hangzhou) unilaterally implanted into the VTA (AP −3.20 mm, ML −0.25 mm, DV −3.8 mm), MD (AP −1.3 mm, ML −0.30 mm, DV −3.2 mm), the SC (AP −3.80 mm, ML ±0.8 mm, DV −1.8 mm), and the BLA (AP −1.5 mm, ML ±3.1 mm, DV −4.70 mm). To block backpropagation of virus in SC, cannulas were implanted 0.3 mm above the SC (AP −3.80 mm, ML ±0.8 mm, DV −1.5 mm). Either bupivacaine (4%, 0.3 μl) or saline (control) was delivered into the SC 30 min before optogenetic modulation and behavioral tests. Mice had at least 2 weeks to recover after surgery before testing.

#### Anatomical tracing

To investigate the origin of the SC-MD and SC-VTA or insula-MD and insula-VTA projecting neurons, we injected RV-dG-dsRed into the VTA and RV-dG-EGFP into the MD in the same animal. To demonstrate the different SC and insula outputs, we injected AAV9-*EF1α*–mCherry into the SC and AAV9-*EF1α*–EYFP into the insula. To map the axonal output of input-defined neurons in the VTA and MD, anterograde *trans*-synaptic AAV1-hSyn-SV40 NLS-Cre was unilaterally injected into the SC, *EF1α*:: DIO–mCherry was injected into the VTA and *EF1α*:: DIO–EYFP was injected into MD.

#### Histology

Mice were given an overdose of pentobarbital and perfused with 0.9% saline followed by 4% paraformaldehyde (PFA) in PBS. Brains were dissected and postfixed in 4% PFA at 4 °C for 24 h then transferred to 30% sucrose for 2 d. Coronal slices (40 μm) were taken across the entire rostrocaudal extent of the brain using a cryostat at −15 °C and stored in 24-well plates containing cryoprotectant at 4 °C. To visualize virus expression, optic fiber tips, optrode placements, and viral tracing targets, floating sections were blocked with 10% normal goat serum in PBS-T (0.03% Triton-X 100), and DAPI (1:50000, Cat#62248, Thermofisher). Brain sections were mounted and cover-slipped with Fluoromount aqueous mounting medium (Sigma-Aldrich, USA). Sections were then photographed using an Olympus VS120 virtual microscopy slide scanning system or a Zeiss LSM LSM 880 confocal microscope. Images were analyzed with ImageJ, Image Pro-plus, and Photoshop software.

#### **c-** Fos Immunolabeling

To visualize c-Fos activity across the whole brain following optogenetic activation of specific neural circuits, we used different optogenetic stimulus groups: *CaMKIIa*::ChR2_Insula-BLA_, *CaMKIIa*::mcherry_Insula-BLA_ (control), and *CaMKIIa*::ChR2_Insula-MD_ *CaMKIIa*::mcherry_Insula-MD_ (control). Mice in each group were given 3 min habituation time and 2 presentations of optogenetic stimuli (20 Hz, 5 ms, 2.5 s, with an interval no less than 2 min) in a looming box during a 10–20 min session. Mice were sacrificed 1.5 hours following optogenetic activation and brains then stained for both c-Fos and DAPI. Sections were washed and incubated in primary (1:500, Cat#2250. CST) and secondary (1:300, Cat#111-547-003, Jackson immuno research) antibodies and DAPI (1:50000, Cat#62248, Thermofisher). Imagines were taken using an Olympus VS120 virtual microscopy slide scanning system or a Zeiss LSM LSM 880 confocal microscope and then overlaid with The Mouse Brain in Stereotaxic Coordinates to locate brain nuclei. Then, c-Fos positive neurons were manually counted by an individual experimenter blind to the experiment groups using ImageJ and Photoshop software.

#### Looming test

The looming test was performed in a closed Plexiglas box (40 ⅹ 40 ⅹ 30 cm) with a shelter nest in one corner. An LCD monitor was placed on the ceiling to present looming stimulus to the upper visual field. The stimulus was a black disc on a grey background expanding from a 2°to 20°visual angle, repeated 15 times, lasting a total of 5.5 seconds. Mouse behavior was recorded using a Sony FDR-AX45 camera. Mice were handled and habituated for 10 min to the looming box one day prior to the test. On the test day, mice were given 3 min to habituate to the box, then 10 looming stimulus trials were presented. The stimulus was triggered by the experimenter when the mouse was far from the shelter and the interval between each trial no less than 2 min. For optogenetic activation experiments with looming stimulus, mice received blue light (10 Hz, 473 nm; Aurora-220-473, NEWDOON, Hangzhou) at an intensity of 8 mW at the fiber tips. Light stimulation was delivered 1 s before onset of the looming stimulus and continued until the stimulus ended.

#### Pupillometry

Mice were head-fixed and allowed to run freely on a stationary foam ball with a diameter of 20 cm during testing. Mice were first habituated to the ball for 3 consecutive days before recording began (day 1, 15 min; day 2, 30 min; day 3, 1 hour). One eye of each mouse was illuminated with a 940 nm near-infrared light and the pupillary responses were recorded using a camera (Point Grey, FL3-U3-13E4M, set to 200 fps). Software (LabVIEW) was used to control the camera and process images in real time to obtain pupil data, including x position, y position, diameter of the pupil and the timestamp for each image. To avoid interference between pupil positions and pupil diameter accuracy, the ellipse long diameter was measured instead of the cross diameter and area.

Looming stimuli were presented using Matlab Psychtoolbox and displayed on a 19-inch screen (DELL, P1917S) during pupillometry recording. An area of 60 x 60 pixels in one corner of the screen light up during the test as a way of modulating brightness. Brightness changes were detected using a photodiode and these signals were transmitted to LabVIEW using the same circuit board and used to timestamp the stimuli time with the pupillary data.

#### Optogenetic manipulation

A closed Plexiglas box (40 x 40 x 30 cm) with a shelter/nest in the corner was used for optogenetic stimulation experiments. Animals were handled and habituated to the looming box for 10 min one day prior to testing. During the looming test session, mice were allowed to freely explore the looming box for 3–5 min and then received either optogenetic manipulation or presentation of the looming stimuli. For optogenetic stimulation experiments, the implanted optic fibers were connected to a 473-nm blue light laser (Aurora-220-473, NEWDOON, Hangzhou) at approximately 15-20 mW for terminals stimulation. Optogenetic activation of neural circuits began 4–5 weeks after animals received stereotactic viral injections and fiber implants. During experiments, light was delivered to either insula-BLA terminals, SC−VTA terminals, insula-VTA terminals, SC−MD terminals or insula-MD terminals. The light was delivered into the targeted regions simultaneously 1 s before onset of the looming stimulus and continued until 1 s after the end of the looming stimulus. Mice received 7.5 s blue light stimulation (150 laser pulses of 5 ms at 20 Hz) at axon terminals during pathway activation experiments. Three repeated light stimuli trials were delivered at about 3 min intervals via a manual trigger in the targeting regions and all light stimulation was manually presented by the experimenter. For all gain-of function experiments (optogenetic activation of ChR2), the activation was all unilateral.

#### Behavioral analysis

Behavioral data were analyzed using Adobe Premiere software and observers were blind to experimental conditions. Mice were allowed to move freely in the open field with a shelter/nest before looming stimulus or light stimulation. Individual time courses were represented setting T=0 ms as the time of stimulation. Three parameters extracted from the behavioral experiments were used to quantify the looming-evoked or light-evoked defensive behavior: (1) rearing frequency: frequency of rearing on hindlimbs and leaning against the walls with one or both forepaws were visually monitored in 1-min blocks. (2) response latency: time between the onset of the looming stimulus or photostimulation and the onset of the escape, escape was defined as the motion that resulted in shelter entrance within stimulation period. (2) return time: the time from looming stimulus or photostimulation presentation to time when the mouse entered the nest. (3) Duration in nest: time spent in the nest following looming stimulus or photostimulation. Data obtained from mice with imprecise fiber placements were not used for analyses.

#### In vivo extracellular recording

Mice were habituated to the head-fixed position whilst on a foam ball 1 hr each day for 3 days. Then, using a 16-channel micro-electrode (Neuronexus, A4×4-6mm-100-125-177-A16) and a multi-channel recording system (OmniPlex D, Plexon, Dallas, USA), the target brain regions were recorded. Electrodes were connected to a headstage (Plexon, Dallas, USA) containing 16-32 unity-gain operational amplifiers. The headstage was connected to a 16-channel computer-controlled preamplifier (gain X-100, band-pass fifilter from 150 Hz to 40 kHz, Plexon). Neuronal activity was digitized at 40 kHz and band-pass filtered from 300 Hz to 8 kHz, and isolated by time-amplitude window discrimination and template matching using a Multichannel Acquisition Processor system (Plexon). To investigate optogenetic effects, we inserted an optic fiber (200 µm diameter; 0.22 NA) 50 µm above the stimulated brain region. Before optical stimulation, we recorded for approximately 3 min to establish a stable electrode position inside the brain tissue. When the signal in the BLA was stable, optical stimulation was delivered at 1 min intervals. At least 10 repetitions of light stimulation was recorded during each session. At the conclusion of the experiment, recording sites were marked with DiI cell labeling solution (Invitrogen, USA) before perfusion, and electrode locations were reconstructed using standard histological techniques.

#### Spike sorting

Single-unit spike sorting was performed using Plexon Offline Sorter software (Plexon, Inc., Dallas, TX, USA), then analyzed in Neuroexplorer (Nex Technologies, Madison, AL, USA) and Matlab (MathWorks, Natick, MA, USA). Principal-component scores were calculated for unsorted waveforms and plotted in a three-dimensional principal-component space; clusters containing similar valid waveforms were manually defined. A group of waveforms was considered to be generated from a single neuron if it defined a discrete cluster in principal component space that was distinct from clusters for other units and if it displayed a clear refractory period (>1 ms) in the auto-correlogram histograms. To avoid analysis of the same neuron recorded on different channels, we computed cross-correlation histograms. If a target neuron presented a peak of activity at a time that the reference neuron fired, only one of the two neurons was considered for further analysis.

To determine whether the firing rate of a particular BLA neuron was altered in response to optogenetic activation of the insula (or SC-VTA,SC-MD, insula, insula-MD) axon terminals, we used peristimulus time histograms (PSTHs) to analyze firing pattern (Buzsaki et al., 2004). We calculated PSTHs using a 7.5 second period before and after onset of optogenetic stimulus with a bin size of 100 ms. We calculated the basal spontaneous firing rate of each neuron by averaging the PSTH over the pre-stimulus bins. Peak optogenetically-evoked firing rate was then calculated as the maximum value of the PSTH after stimulus onset (within 2.5 seconds from the stimulus). The baseline mean was the average of the PSTH bins before stimulus onset, and the SD was the standard deviation of the PSTH bins before stimulus onset. We calculated a Z-score firing rate using the following equation: Z = (FR − mean of FRb)/SD of FRb, where FR indicates the firing rate for each bin and FRb indicates the baseline firing rate before the stimulus onset. A positive responding neuron was defined when the absolute value of the Z-score firing rate of least one time bin after stimulation was larger than 2. Negative responding neurons were defined when the absolute value of the Z-score firing rate of least one time bin after stimulation was smaller than –2.

#### Power spectrum analysis

The LFP data before and after optogenetic stimulation was conducted with power spectrum analysis, which is a technique for decomposing complex signals into simpler signals based on Fourier transform. The power spectral density (PSD) for LFP data was computed using the multi-taper method (TW = 3, K = 5 tapers) using the Chronux toolbox using custom-written or existing functions in MATLAB (The Math Works).

#### Statistical analysis

The number of biological replicates in each group was 3–6 mice per group for anatomy, 7–9 mice for optogenetic manipulation and looming stimulation, and 16–51 mice per group for behavioral habituation to looming stimulus. Data distribution was assumed to be normal, but this was not formally tested. All statistics were performed using Graph Pad Prism (GraphPad Software, Inc.) and MATLAB. Paired student T-test, unpaired student T-test, one-way ANOVA and two-way ANOVA were used where appropriate. Bonferroni post-hoc comparisons were conducted to detect significant main effects or interactions. In all statistical measures, a P value <0.05 was considered statistically significant. Post-hoc significance values were set at *P< 0.05, **P< 0.01, ***P< 0.001 and ****P< 0.0001, and all statistical tests used are indicated in the figure legends.

## Figures and figure legends

**Supplementary Figure 1.**
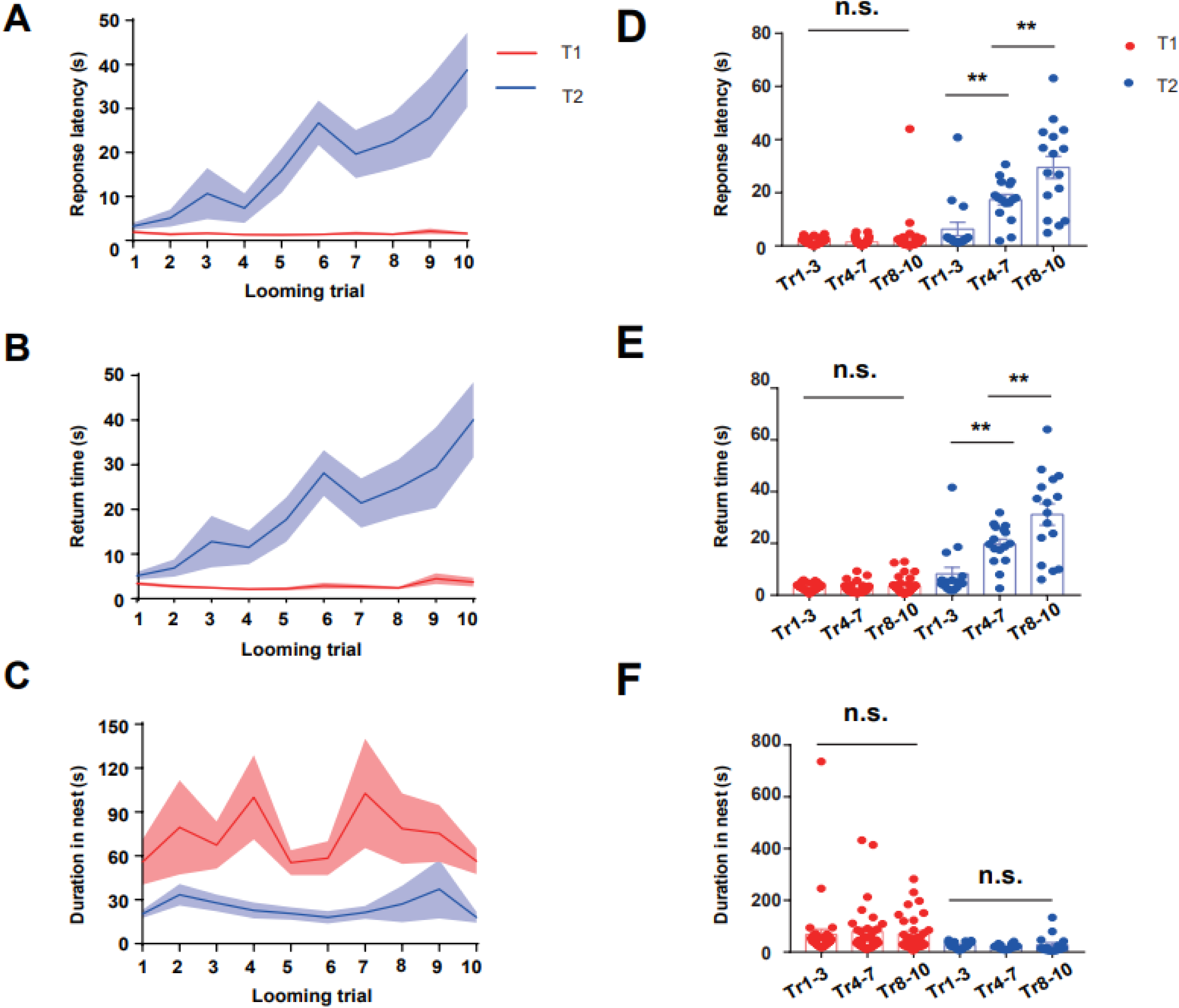
Behavioral Characteristics of T1 and T2 Groups. **(A)** Graph showing the average response latency in T1 and T2 mice. **(B)** Graph showing the average return time in T1 and T2 mice. **(C)** Graph showing the average time spent in nest in T1 and T2 mice. **(D)** Bar graph showing the average response latency in trials 1–3, trials 4–7 and trials 8–10 (one-way ANOVA, p>0.05, Two-tailed paired t test, **p=0.0013, Two-tailed paired t test, **p=0.0074). **(E)** Bar graph showing the average return time in trials 1–3, 4–7 and 8–10 (one-way ANOVA, p>0.05, two-tailed unpaired t test, **p=0.0011, two-tailed unpaired t test, **p=0.01). **(F)** Bar graph showing the average duration in the nest during trials 1–3, 4–7 and 8–10 (one-way ANOVA, p>0.05, two-tailed unpaired t test, p>0.05, two-tailed unpaired t test, p>0.05) of T1 and T2 mice responding to 10 blocks of LS.

**Supplementary Figure 2.**
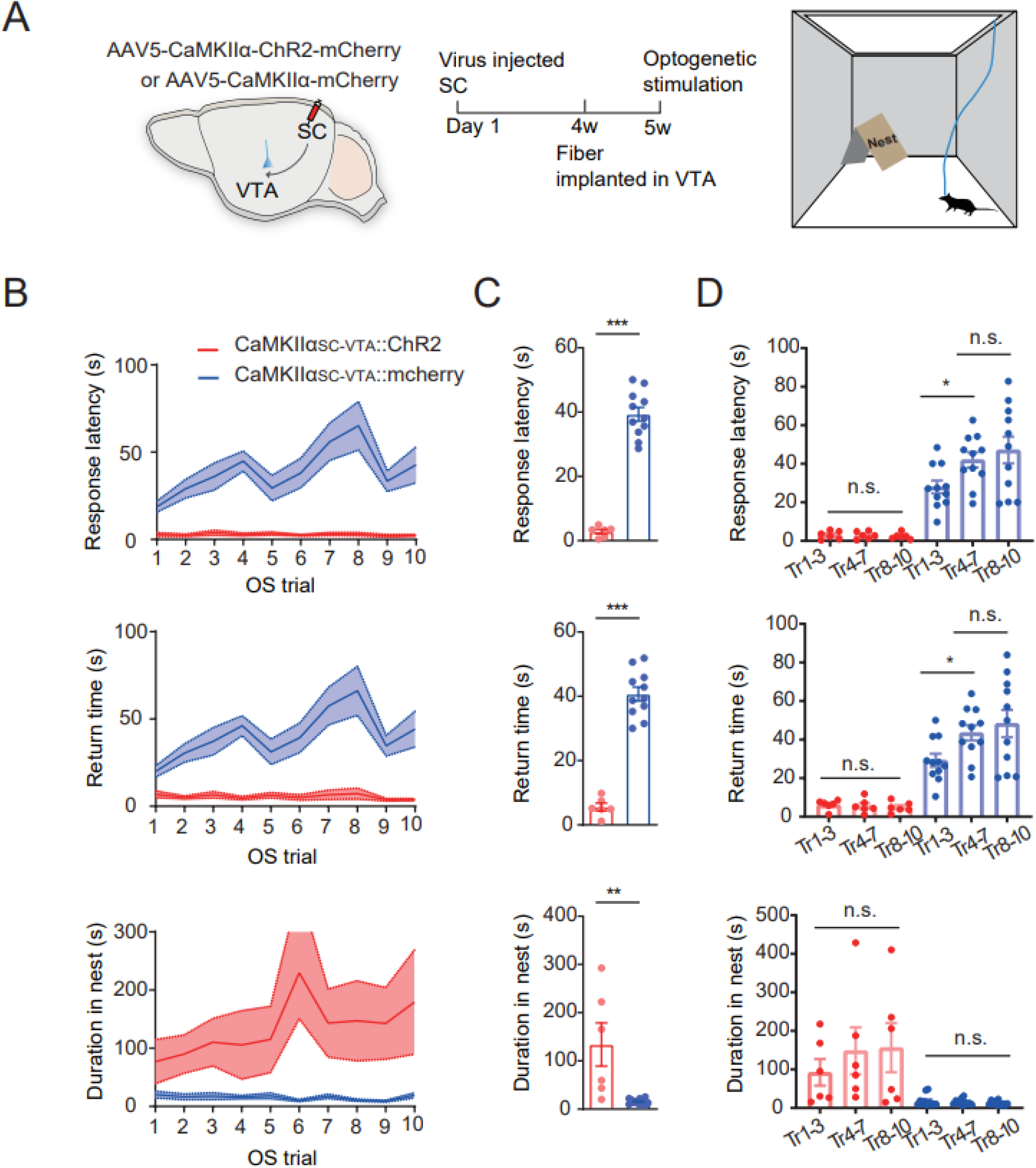
Selective repeated activation SC-VTA mimicked T1-type behaviors. (**A**) Schematic showing the injection of AAV-CaMKIIα-ChR2-mcherry into the IDSC and the experimental timeline and optogenetic stimulation (OS) paradigm. (**B**) Time course analysis of 1) response latency, 2) return time and 3) duration in nest following optogenetic stimulation of both the CaMKIIα_SC-VTA_:: ChR2 and mcherry control groups. (**C**) Measurements were taken during OS (10 trials/mouse) of the CaMKIIα_SC-VTA_:: ChR2 and mcherry groups. Bar graph showing the average 1) response latency (Two-tailed t test, **** p <0.0001), 2) return time (two-tailed t test, **** p<0.0001) and 3) duration in nest (two-tailed t test, **p=0.0023). (**D**) Measurements were taken during OS (10 trials/mouse) of the CaMKIIαSC-VTA:: ChR2 and mcherry groups. Bar graphs showing average values in trials 1–3, 4–7, and 8–10 for 1) response latency (one-way ANOVA, p=0.6764, *p=0.0296), 2) return time (one-way ANOVA, p=0.3136,*p=0.0325) and 3) duration in nest (one-way ANOVA, p=0.4208,p=0.4858).

**Supplementary Figure 3.**
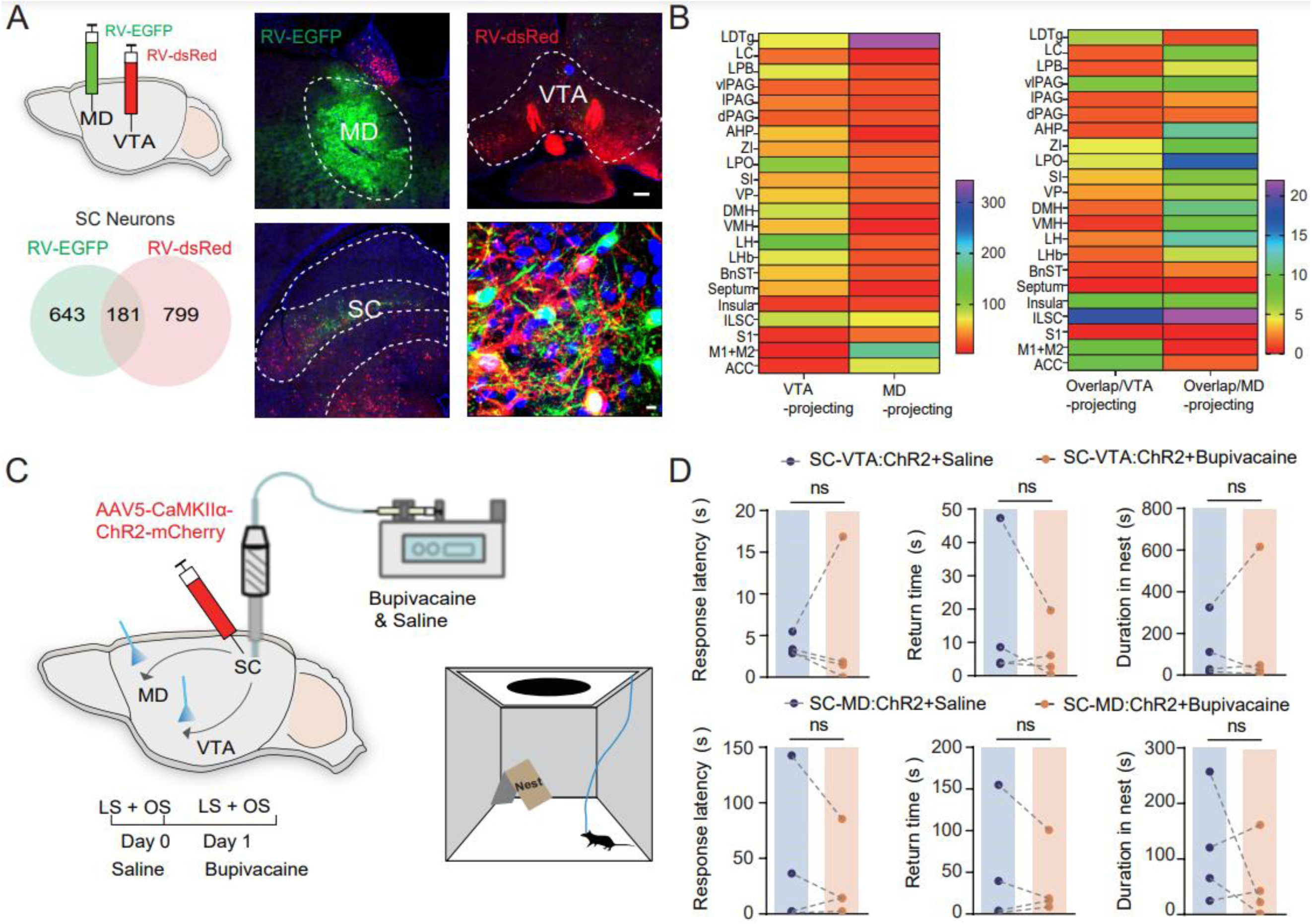
SC Pathway Parallel Processing Modulates Variability in Innate Escape Responses. **(A)** Schematic showing injections of RV-dsRed and RV-EGFP into VTA, the MD and the number of SC neurons labeled by each, and number that overlapped. **(B)** Heatmap showing quantification of the number of RV-dsRed-and RV-EGFP-labeled neurons, and the amount of overlap, across the whole brain. **(C)** Schematic showing the blocking of neuronal backpropagation in the SC and the experimental timeline. **(D)** Bupivacaine injection (0.3 μl 4%) in the SC had no effect on stimulation of the SC-VTA and SC-MD. Measurements were taken during OS (10 trials/mouse) and LS of the CaMKIIαSC-VTA:: ChR2 and the CaMKIIα_SC-MD_:: ChR2 groups given either saline or bupivacaine. Average values in trials 1–3, 4–7, and 8–10 were calculated. *Top* Bar graphs for the CaMKIIαSC-VTA:: ChR2 group showing 1) response latency (two-tailed t test, p>0.05), 2) return time (two-tailed t test, p>0.05) and 3) duration in nest (two-tailed t test, p>0.05). *Bottom* Bar graphs for the CaMKIIα_SC-MD_:: ChR2 group showing 1) response latency (two-tailed t test, p>0.05), 2) return time (two-tailed t test, p>0.05) and 3) duration in nest (two-tailed t test, p>0.05).

**Supplementary Figure 4.**
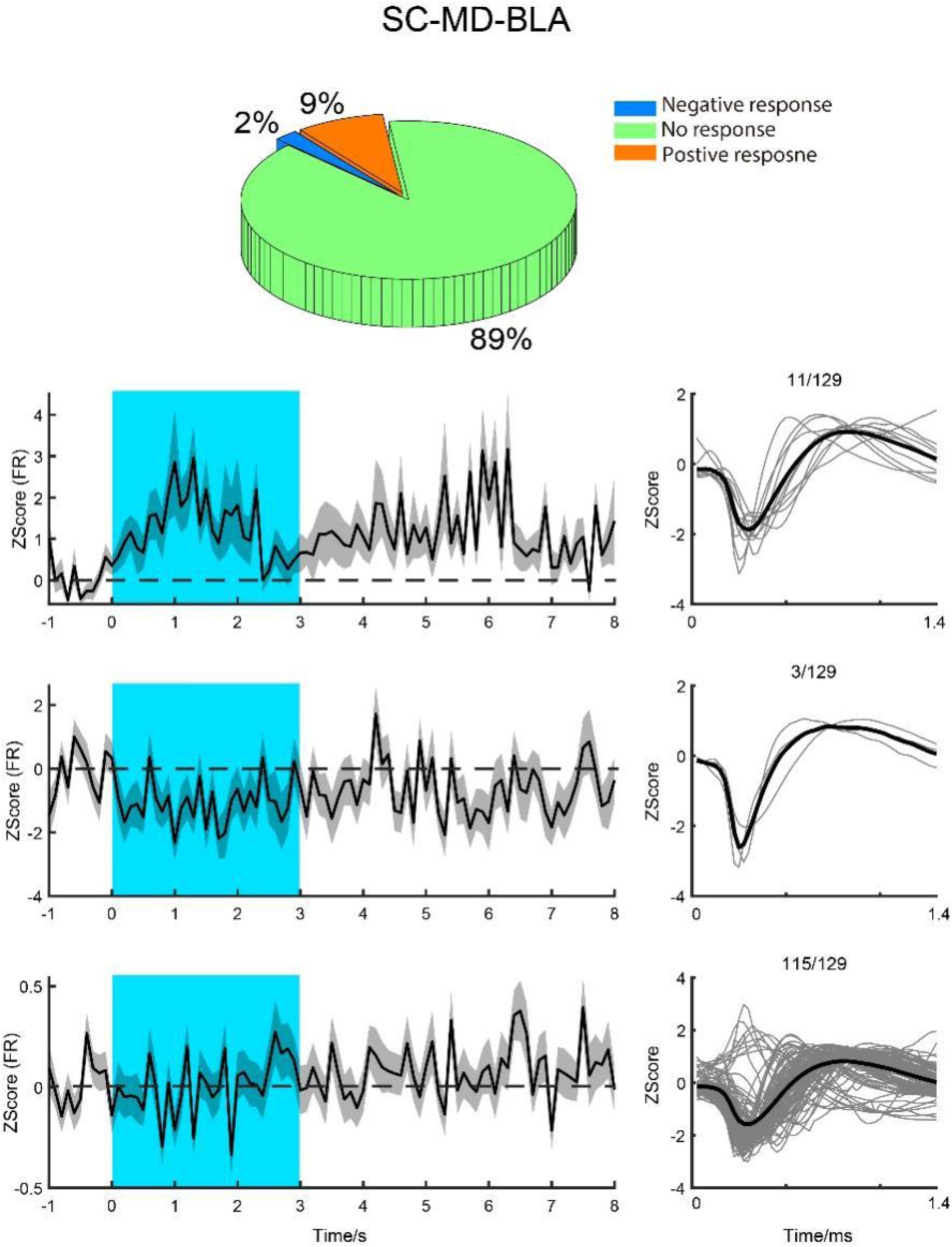
BLA neuronal responses to activation of the SC-MD pathway. Pie graph showing the proportion of neurons recorded in BLA with a positive (orange), negative (blue) and no response (green) to optical stimulation of the SC-MD pathway. Line graphs showing the firing rates (z-scores) of three neuronal types before, during and after optogenetic stimulation (blue shadow) alongside the corresponding spike waveforms (*right*).

**Supplementary Figure 5.**
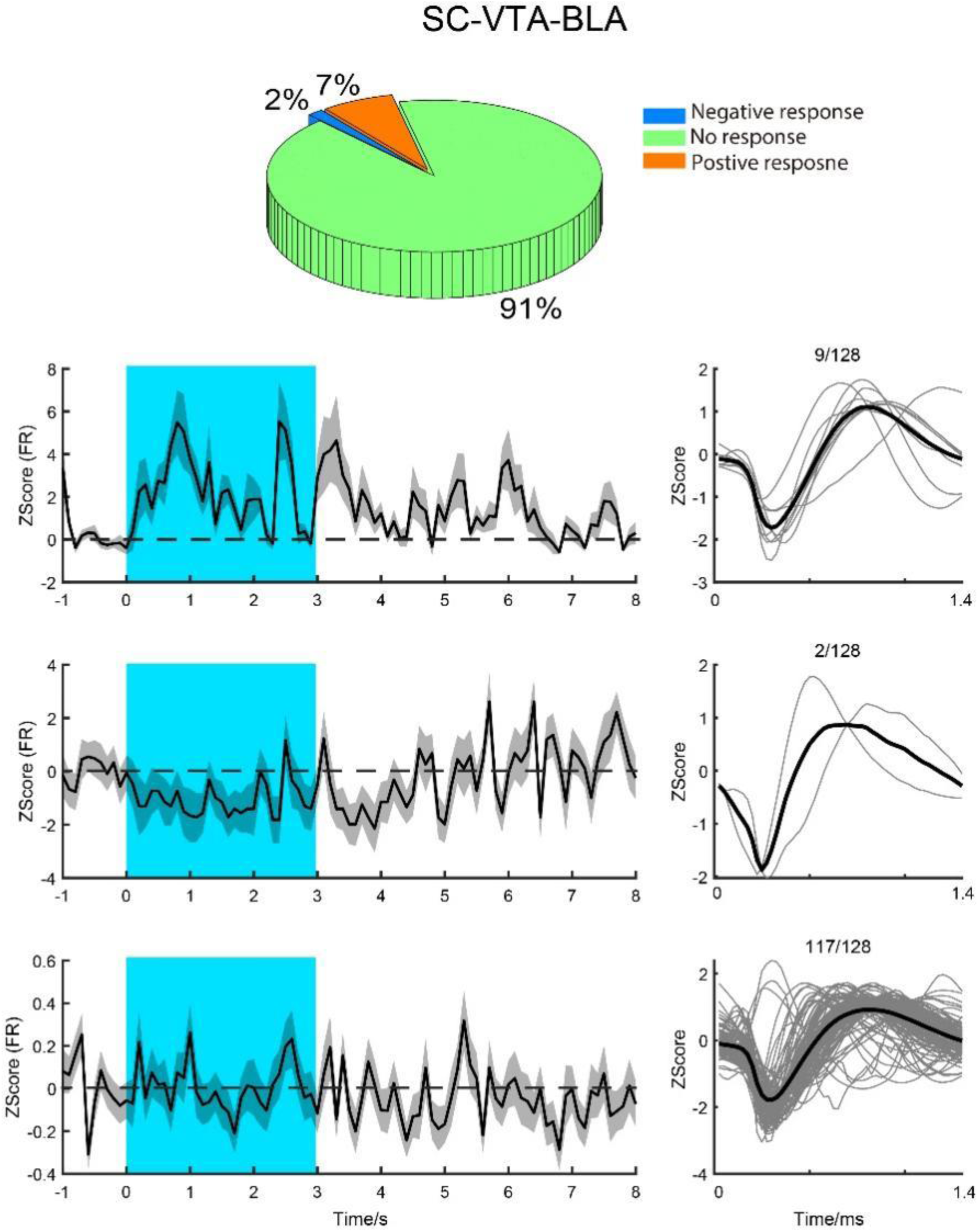
BLA neuronal responses to activation of the SC-VTA pathway. Pie graph showing the proportion of neurons recorded in the BLA with positive (orange), negative (blue) and no response (green) to optical stimulation of the SC-VTA pathway. Line graphs showing the firing rates (z-scores) of three neuronal types before, during and after optogenetic stimulation (blue shadow) alongside the corresponding spike waveforms (*right*).

**Supplementary Figure 6.**
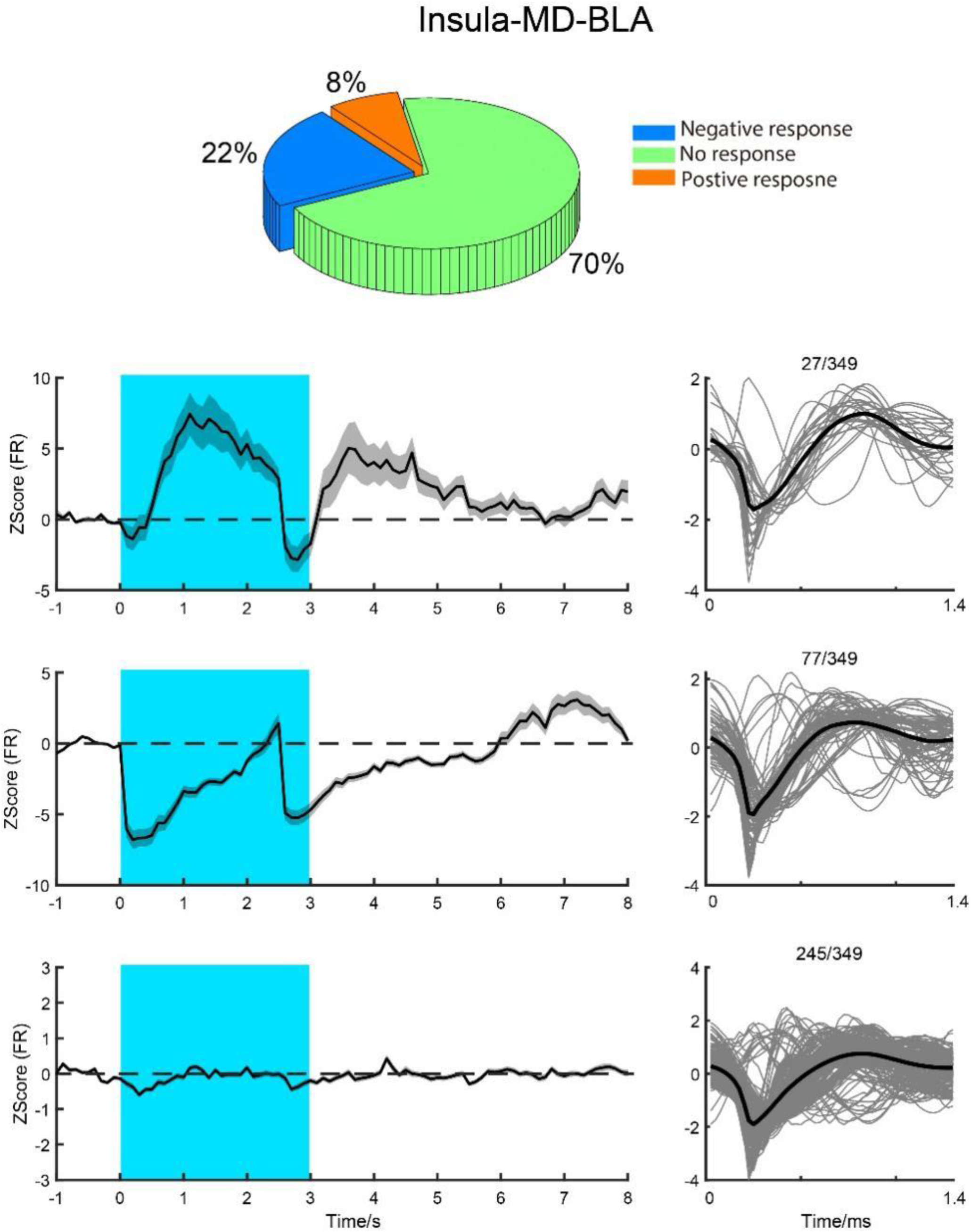
BLA neuronal responses to activation of the Insula-MD pathway. Pie graph showing the proportion of neurons recorded in the BLA with positive (orange), negative (blue) and no response (green) to OS of the insula-MD pathway. Line graphs showing the firing rates (z-scores) of three neuronal types before, during and after optogenetic stimulation (blue shadow) alongside the corresponding spike waveforms (right).

**Supplementary Figure 7.**
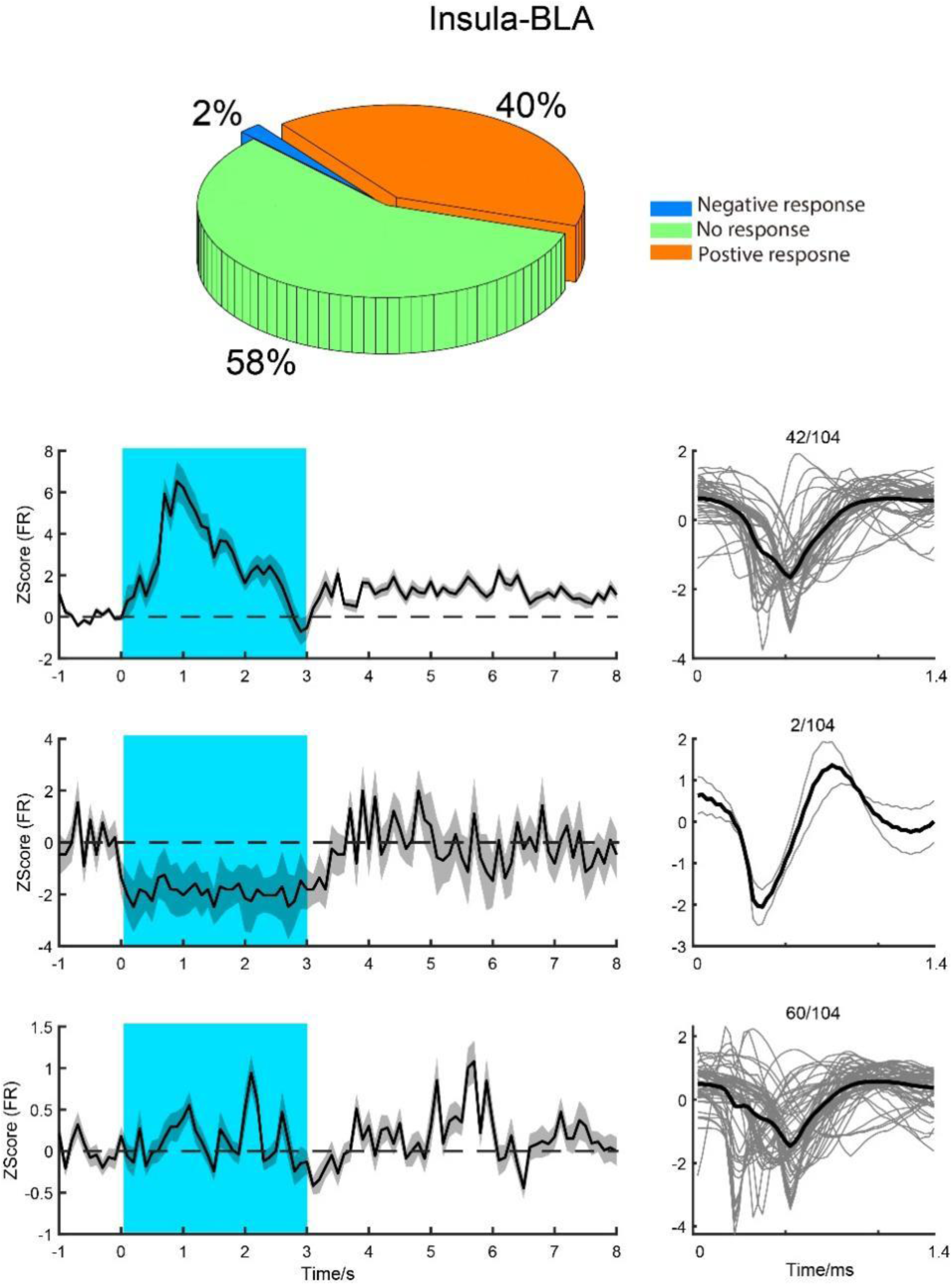
BLA neuronal responses to activation of the Insula-BLA pathway. Pie graph showing the proportion of neurons recorded in BLA with positive (orange), negative (blue) and no response (green) to OS. Line graphs showing the firing rates (z-scores) of three neuronal types before, during and after optogenetic stimulation (blue shadow) alongside the corresponding spike waveforms (right).

