## Supplementary figures and images for "Neural Circuit Underlying Individual differences in Visual Escape Habituation"

### Supplemental Figure 1

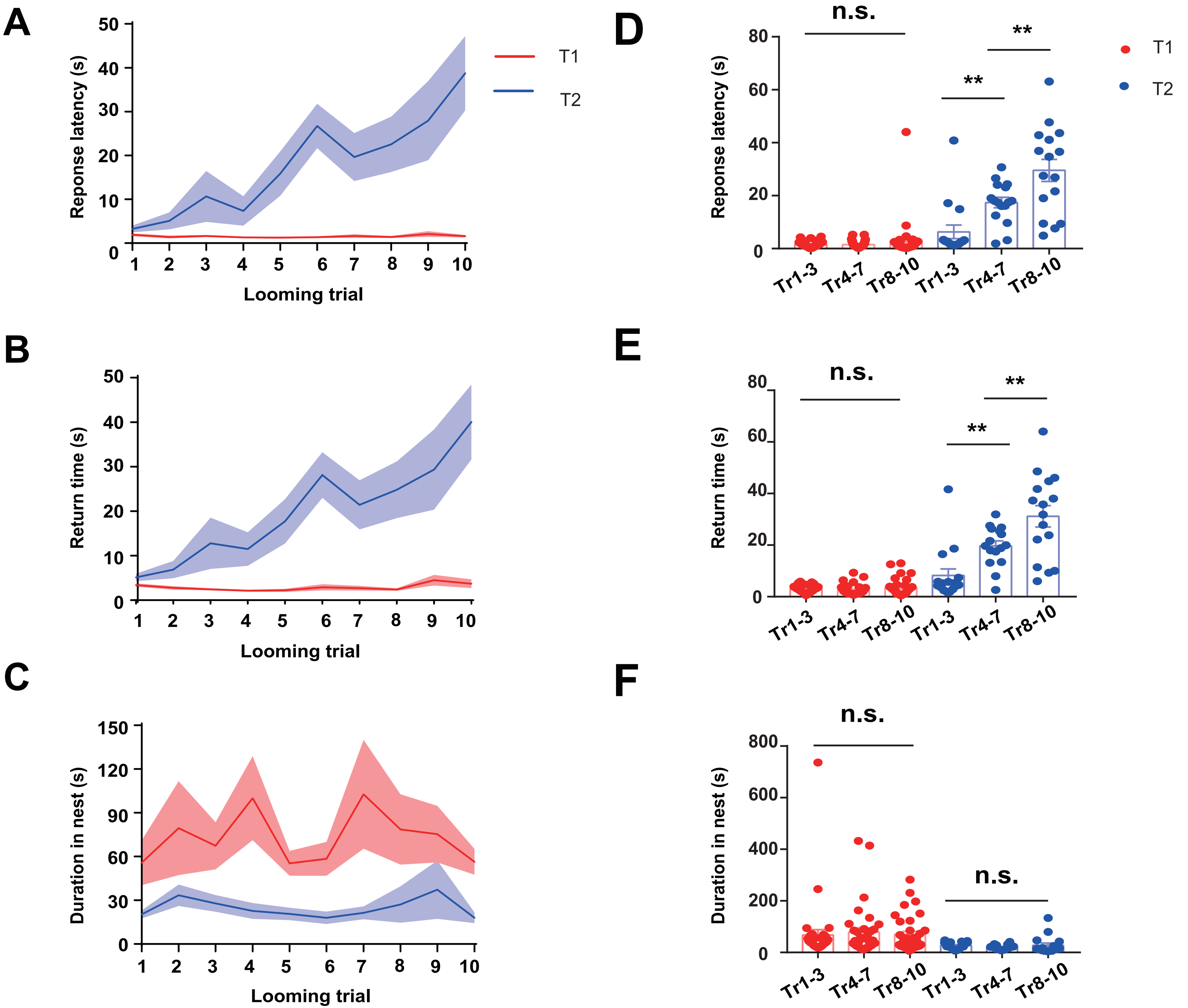

### Supplemental Figure 2

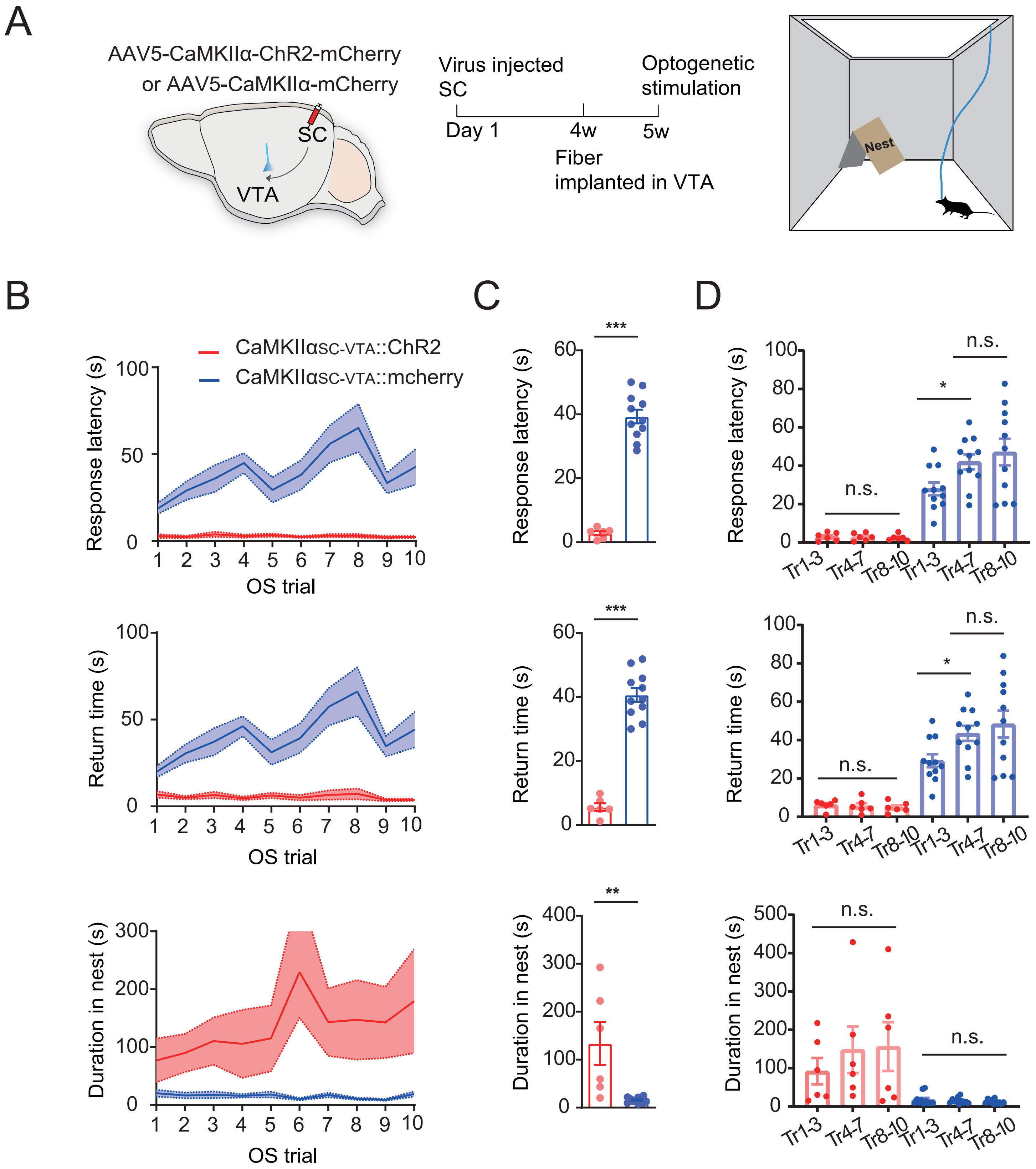

### Supplemental Figure 3

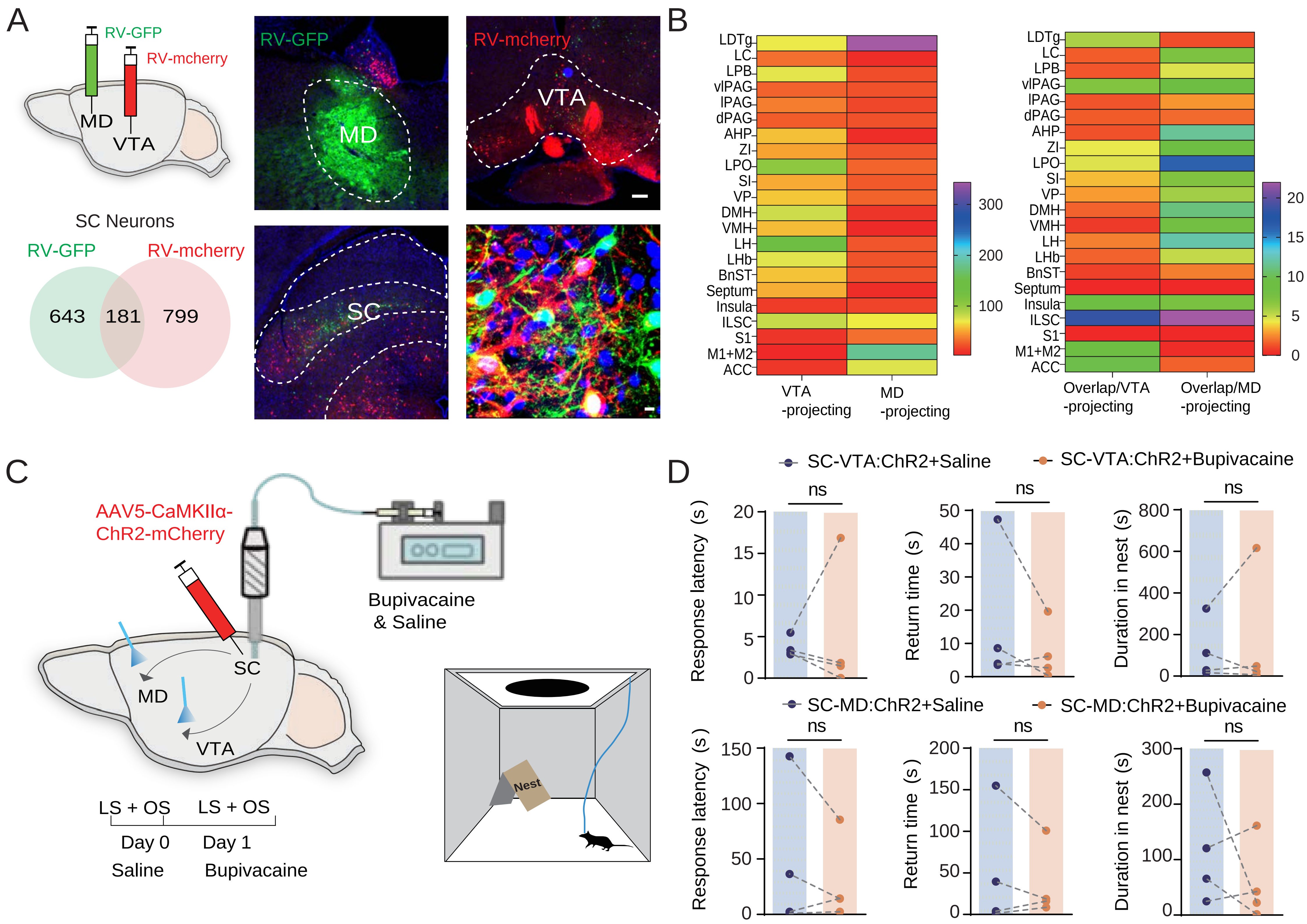

### Supplemental Figure 4

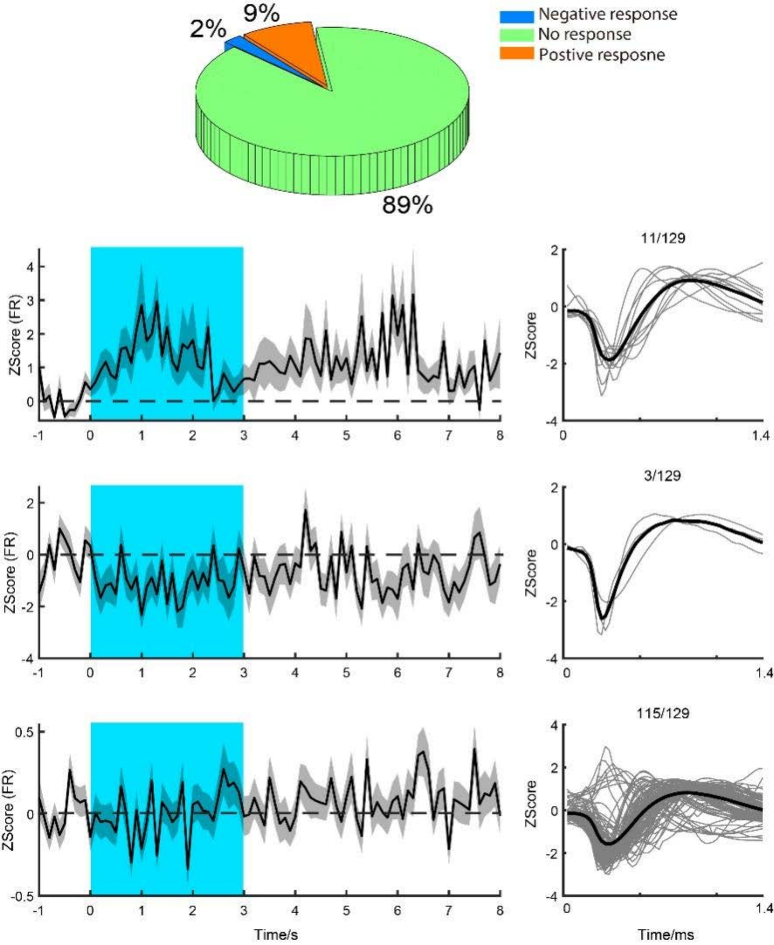

### Supplemental Figure 5

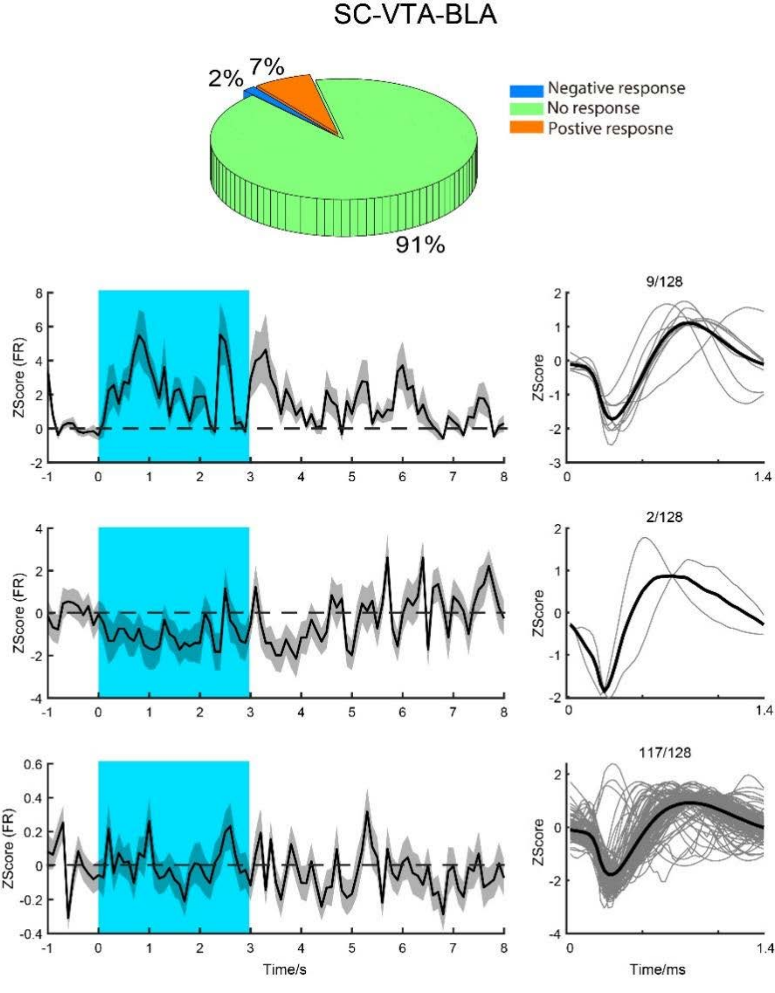

### Supplemental Figure 6

#### Insula-MD-BLA

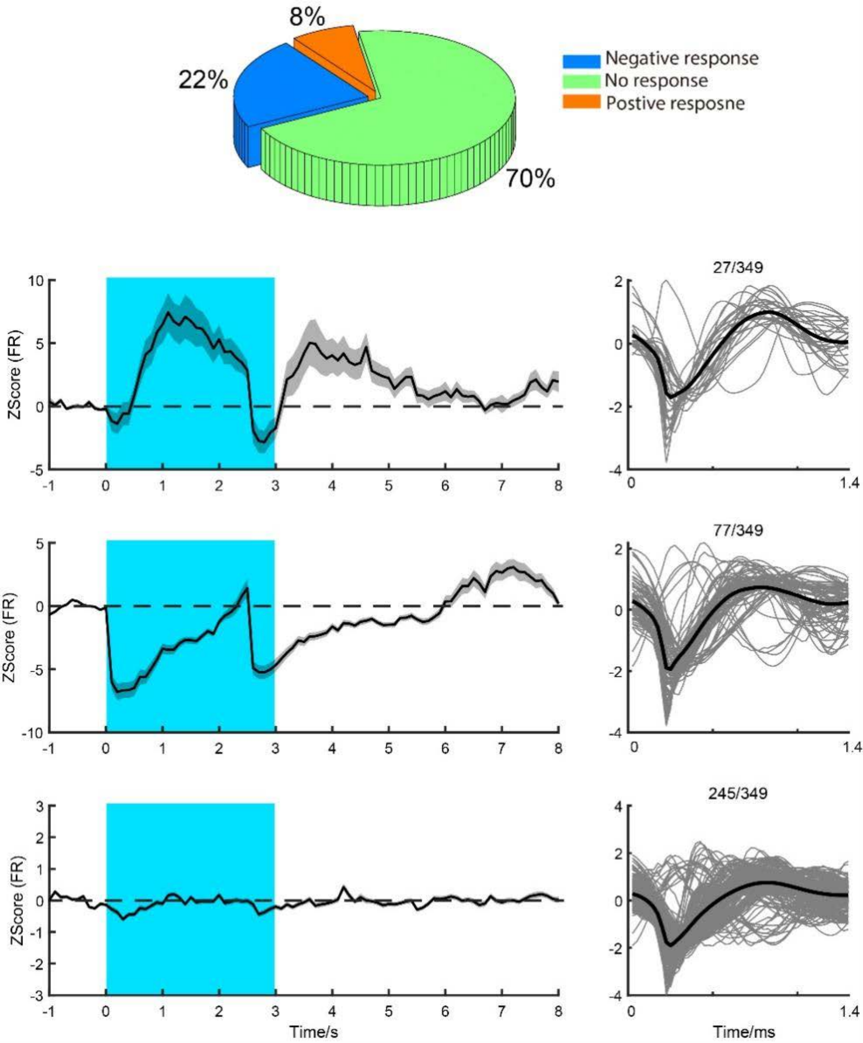

### Supplemental Figure 7

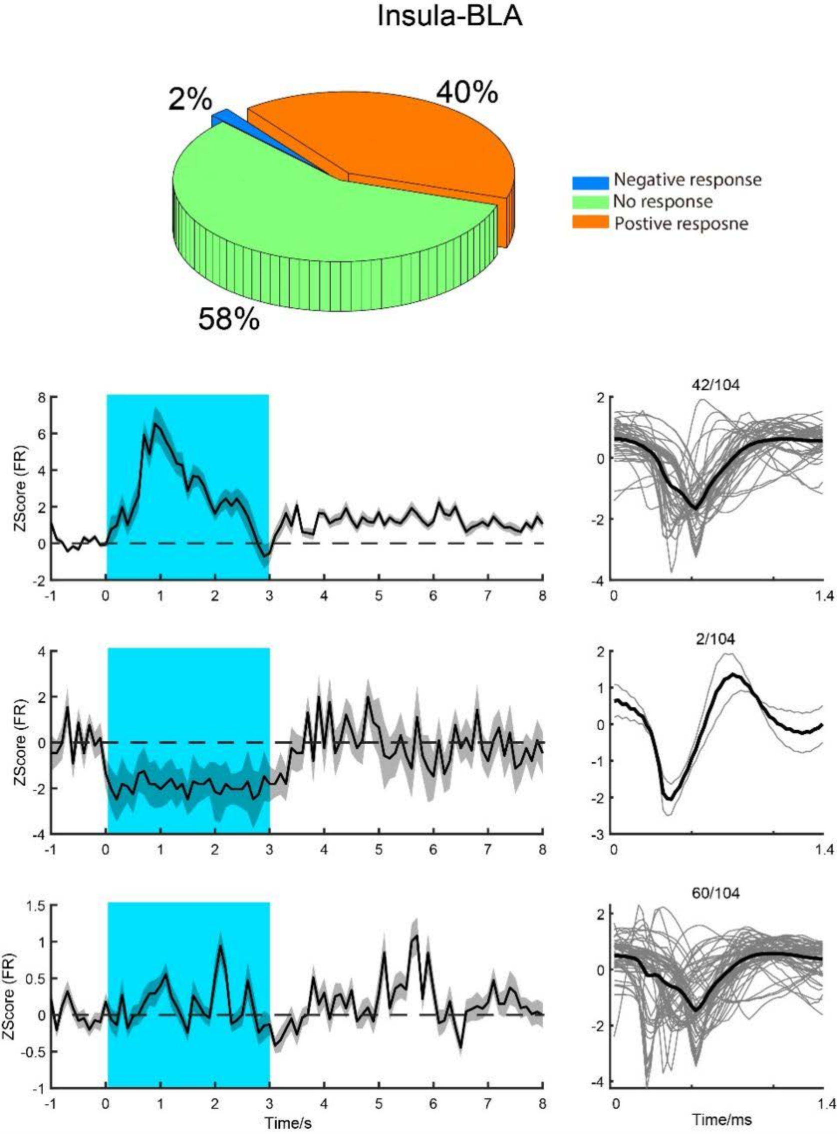
